# Erythroid differentiation intensifies RNA mis-splicing in *SF3B1*-mutant myelodysplastic syndromes with ring sideroblasts

**DOI:** 10.1101/2023.04.11.536355

**Authors:** Pedro L. Moura, Teresa Mortera-Blanco, Isabel J.F. Hofman, Gabriele Todisco, Warren W. Kretzschmar, Ann-Charlotte Björklund, Maria Creignou, Michael Hagemann-Jensen, Christoph Ziegenhain, David C. Granados, Indira Barbosa, Gunilla Walldin, Monika Jansson, Neil Ashley, Adam J. Mead, Vanessa Lundin, Marios Dimitriou, Tetsuichi Yoshizato, Petter S. Woll, Seishi Ogawa, Rickard Sandberg, Sten Eirik W. Jacobsen, Eva Hellström-Lindberg

## Abstract

Myelodysplastic syndromes with ring sideroblasts (MDS-RS) commonly originate from mutations in the splicing factor *SF3B1* (*SF3B1*^mt^). *SF3B1*^mt^ cause RNA mis-splicing, mechanistically established as the major driver of RS development. However, little is known about RS fate and biology after their initial formation in the human bone marrow. We here achieve isolation of viable RS from patient samples, enabling the first complete investigation of *SF3B1*^mt^ development from stem cell to RS. We show that RS skew MACS-isolated CD34^+^ data towards erythroid features not recapitulated in single-cell RNAseq. We demonstrate that RS divide, differentiate, enucleate and actively respond to mis-splicing/oxidative stress, decreasing wildtype stem cell fitness via GDF15 overproduction. We identify circulating RS as a uniform clinical feature associated with disease burden. Finally, we establish that *SF3B1*^mt^ mis-splicing intensifies during erythroid differentiation and demonstrate through combined transcriptomics/proteomics an uncoupling of RNA/protein biology in RS encompassing severe and dysfunctional mis-splicing of proapoptotic genes.

**Statement of significance:** We here combine a novel method for RS isolation with state-of-the-art multiomics to perform the first complete investigation of *SF3B1*^mt^ MDS-RS hematopoiesis from stem cell to RS. We identify the survival mechanisms underlying *SF3B1*^mt^ erythropoiesis and establish an active role for erythroid differentiation and RS themselves in *SF3B1*^mt^ MDS-RS pathogenesis.

## Introduction

Myelodysplastic syndromes (MDS) are clonal myeloid malignancies in which one or multiple branches of hematopoiesis are disrupted due to ineffective differentiation of hematopoietic stem and progenitor cells (HSPCs), generally resulting in cytopenia.^2^ Myelodysplastic syndromes with ring sideroblasts (MDS-RS) are a slowly progressing low-risk MDS subgroup comprising 16-24% of all MDS.^3^ MDS-RS primarily affect the erythroid lineage and are characterized by extensive dyserythropoiesis and perinuclear accumulation of mitochondria loaded with aberrant ferritin-iron complexes, reflected as the ring sideroblast phenotype (RS).^4^

Hematopoietic stem cell-borne mutations in the *SF3B1* gene (pre-mRNA-splicing factor 3b subunit 1) are the primary disease driver in more than 80% of MDS-RS cases^5,6^. Moreover, 80% of *SF3B1*^mt^ MDS-RS patients display none or few co-mutations, confirming *SF3B1*^mt^ as the disease-driving molecular event.^7^ *SF3B1* mutations cause extensive RNA mis-splicing,^8,9^ leading to increased nonsense-mediated decay (NMD) of mis-spliced transcripts. Mis-splicing and downstream NMD of *ABCB7* and *PPOX* transcripts in particular is mechanistically associated with RS development by disrupting iron processing^10,11^, which cascades into redox imbalance, obstructed differentiation and increased apoptosis of erythroid precursors.^12,13^

Despite alternative splicing (AS) and NMD supposedly reducing the survivability of *SF3B1*^mt^ cells (particularly erythroid cells), the MDS-RS bone marrow (BM) is hyperplastic and usually displays increased erythropoiesis with abundant RS. The molecular mechanisms that enable RS survival, expansion and accumulation in the MDS-RS BM therefore remain unclear.

Exploring how *SF3B1*^mt^ affects erythropoiesis has been pursued through diverse model systems,^6,14-17^, each of which unable to recapitulate full BM/erythroid biology and/or RS development. Crucially, the isolation of viable human RS has not yet been achieved, complicating direct studies into MDS-RS pathobiology.

By exploiting the accumulation of ferric iron inherent to the RS phenotype, we here isolate RS and investigate the entire course of *SF3B1*^mt^ erythropoiesis through an integrative multiomic approach. We identify several molecular pathways engaged to minimize the consequences of widespread mis-splicing and extreme oxidative stress, promoting RS survival and accumulation. Finally, we identify increased RNA mis-splicing, decreased NMD, and transcriptome-proteome uncoupling during erythroid differentiation which altogether demonstrate a role for *SF3B1*^mt^ erythropoiesis itself in driving disease pathogenesis.

## Results

### SF3B1^mt^ RS remain active during MDS-RS erythropoiesis

To investigate RS development during erythropoiesis, we first pursued erythroblast (EB) staging in mononuclear cells (MNC) from MDS-RS and healthy BM samples through co-detection of Band 3 and Integrin α4^18^ (**Fig. 1A**). MDS-RS samples displayed a clear increase in total erythroid frequency, but with relative accumulation at early precursor stages (Pro/Basophilic EB) and matching depletion at the Orthochromatic EB stage (**Fig. 1B**). Morphological analysis identified a progressive increase in RS frequency with erythroid maturation (**Fig. 1C, Fig. S1-S3**). Assessing mitotic activity showed significantly increased DNA in MDS with no change in Ki-67 expression, indicating uncoupled cell cycle dynamics corresponding to a dysplastic phenotype (**Fig. 1D**). While RS decreased with increasing Ki-67 signal, the median Ki-67 population had equal RS frequencies to diagnostic BM smears (**Fig. 1E**). Interestingly, these experiments also identified aberrantly decreased CD71 surface expression by MDS precursors as a potential mechanism to avoid iron overload (**Fig. 1F**). Finally, erythroid enucleation ability was assessed through 3D cell culture of MDS-RS cells.^15^ RS were found to enucleate at equal rates to normoblasts from the same patients (**Fig. 1G, Fig. S4**), fully recapitulating progression through erythropoiesis.

**Figure 1:**
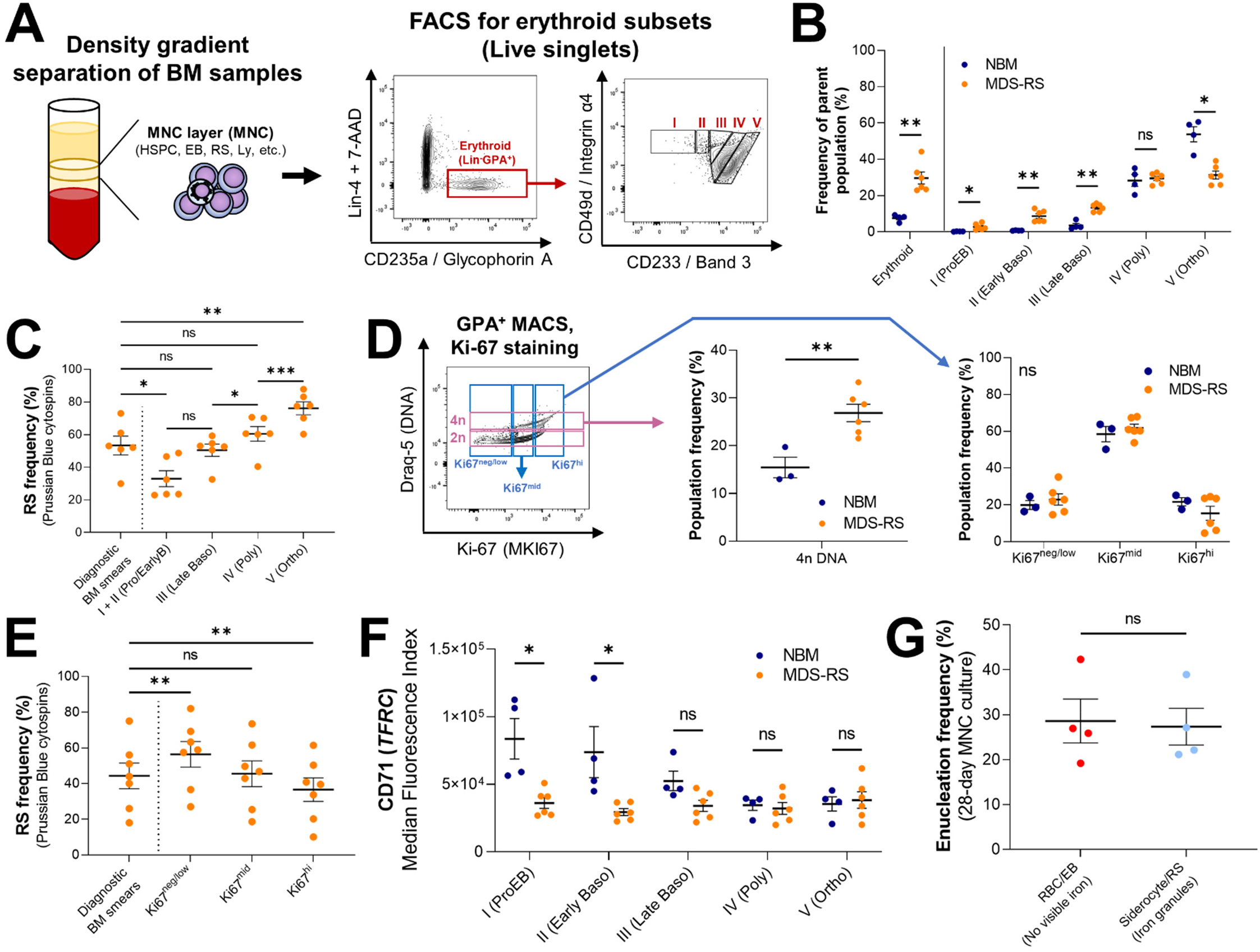
Erythroid differentiation and enucleation remain active in *SF3B1*^mt^ MDS-RS erythroblasts. **A)** Flow cytometry (FC) strategy for staging of erythroblast populations from bone marrow (BM) mononuclear cells (MNC) of patients with myelodysplastic syndromes with ring sideroblasts (MDS-RS). Gating steps identify live and terminally differentiating erythroid cells (Lin^-^7AAD^-^ GPA^+^), from which erythroblasts (EB) are staged according to Band 3 and Integrin α4 expression.^18^ Integrin α4-negative cells are excluded from quantification to avoid skewing by anucleate cells. **B)** Mean (± standard error of the mean, SEM) cell population frequencies within FC parent populations (singlets > terminally differentiating erythroid > EB subsets), quantified in normal bone marrow (NBM) donors (n^NBM^ = 4) and MDS-RS patients (n^MDS-RS^ = 6). Erythroid cells are quantified within the singlet population, EB subsets are quantified within the GPA^+^ population. **C)** Mean (± SEM) ring sideroblast (RS) frequencies per sorted EB subset and compared with frequencies in matched diagnostic BM smears (n^MDS-RS^ = 6). **D)** Gating and quantification [Mean (± SEM)] of DNA content (Draq-5) and intracellular Ki-67 abundance in GPA^+^ magnetically-sorted cells (n^NBM^ = 3, n^MDS-RS^ = 6). **E)** Mean (± SEM) RS frequencies per sorted Ki67-expressing subset and compared with frequencies in matched diagnostic BM smears (n^MDS-RS^ = 6). **F)** Mean (± SEM) CD71 (transferrin receptor, *TFRC*) median fluorescence indices (MFI) per EB subset (n^NBM^ = 4, n^MDS-RS^ = 6). **G)** Mean (± SEM) enucleation frequencies after 28-day 3D culture of MDS-RS BM MNCs (n^MDS-RS^ = 4) and separated by iron granule visibility upon morphological analysis. Statistical comparison was performed by paired T-test analysis. * = p < 0.05, ** = p < 0.01, *** = p < 0.001, ns = non statistically significant.

### Successful separation of viable SF3B1^mt^ RS/siderocytes enables focused investigations

Further assessment of RS biology was challenging in heterogeneous cell populations. However, as hemozoin-rich red blood cells (RBC) are commonly separated by reagent-free magnetic isolation (MACS),^19^ we hypothesized that iron burden could similarly be exploited to isolate *SF3B1*^mt^ RS/siderocytes (**Fig. 2A**). MACS was performed on MNC (viably frozen) and high-density (HD, fresh) fractions and achieved significant RS enrichment in both (**Fig. 2B**). Fluorescence-activated cell sorting (FACS) of HD-isolated cells further enriched a high-purity (99.3%) RS population at significantly higher abundance than MNC (**Fig. 2C, Fig. S5**) and was the method utilized throughout this study unless otherwise specified. Isolated RS significantly correlated with morphological BM RS frequencies (**Fig. 2D**) and were confirmed as fully clonally involved (**Fig. 2E**). Validating our previous unbiased approaches, purified RS displayed a highly significant decrease in surface CD71 levels (**Fig. 2F, Fig. S6**) and also presented a spectrum of Ki-67 nuclear localization (**Fig. 2G, Fig. S7**).

**Figure 2:**
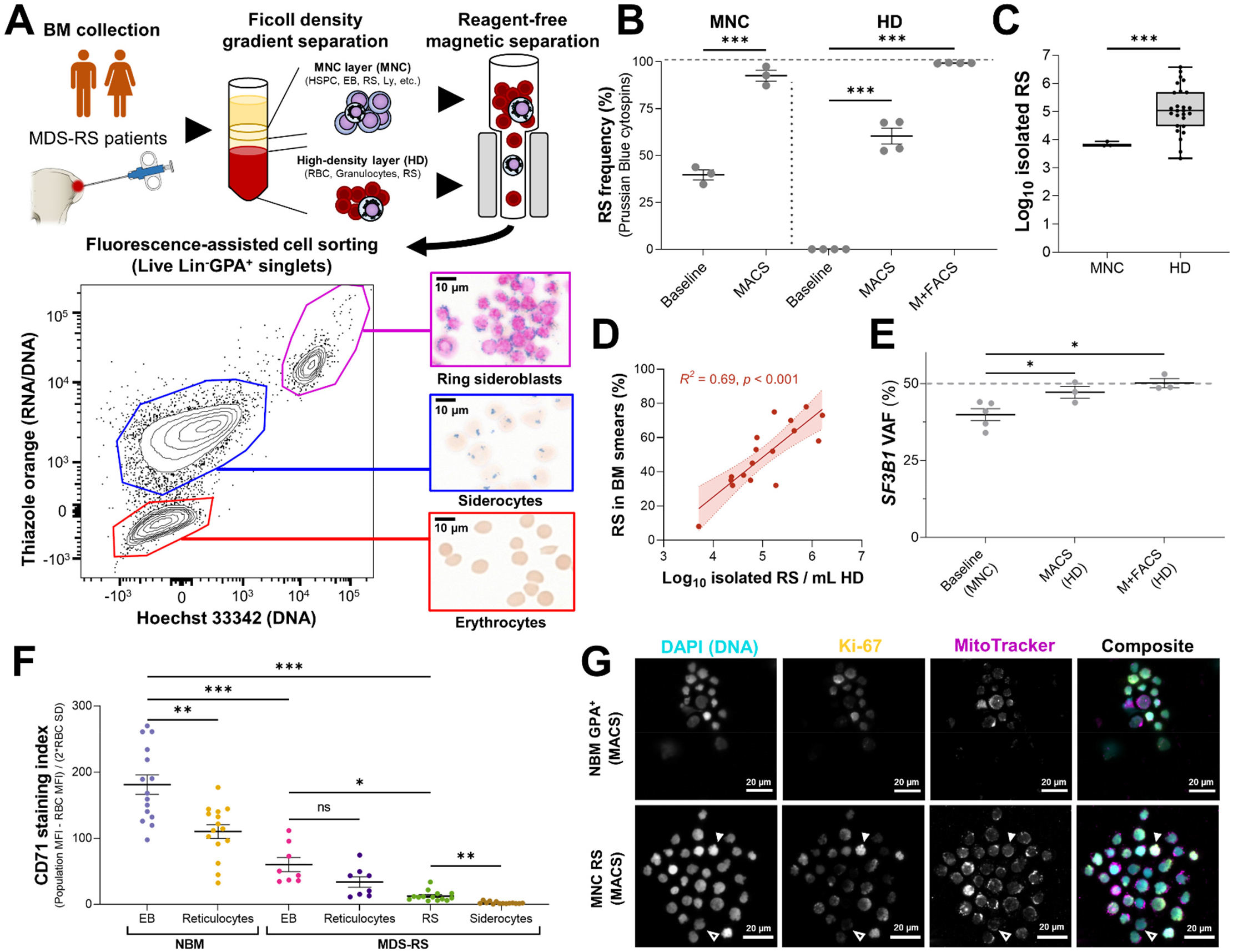
Reagent-free MACS enables direct characterization of viable *SF3B1*^mt^ RS. **A)** Method for RS and siderocyte purification from BM aspiration material. A representative FC diagram plots RNA/DNA content (Thiazole Orange [TO]) against DNA content (Hoechst 33342) in Lin^-^GPA^+^ singlets after MACS of HD cells. Representative micrographs are shown to the right (iron granules in blue, hemoglobin brown and DNA pink). Scale bars: 10 μm. **B** Mean (± SEM) RS frequencies before and after MACS alone in 3 MNC and 4 HD samples, and further purification with FACS (M+FACS) of the same HD samples. HD RS quantification before enrichment steps identifies only 0.1-0.001% as potential RS due to high RBC proportions. **C)** Isolated RS numbers in MACS-enriched cells from 5×10^6^ MNCs (n = 3) or M+FACS-enriched HD cells (n = 26, 19 unique biological replicates + 7 repeat visits). **D)** Correlation of log_10_-converted isolated RS numbers and RS frequencies in matched BM aspirates (n = 17). **E)** Mean (± SEM) *SF3B1* mutation (*SF3B1*^mt^) variant allele frequency (VAF) in unfractionated MNCs (baseline) and MACS-enriched or M+FACS-enriched HD cells, as determined by droplet digital PCR (n = 3 per enrichment method, 5 patients in total). The dashed line indicates complete heterozygosity (VAF = 50%). **F)** Mean (± SEM) CD71 staining indices (MFI of the cell population – MFI of the CD71-negative red blood cell (RBC) population divided by 2 x standard deviation [SD] of the RBC population) (n^NBM^ = 15, n^MDS-RS^ = 8, n^RS^ = 14). **G)** Immunofluorescence of Ki-67 detection in NBM EB and an MNC-derived RS isolate, co-labeled for DNA (DAPI; cyan), Ki-67 (yellow) and mitochondria (MitoTracker; magenta). Individual greyscale channels and a composite image of all three markers are shown. A Ki-67^neg^ RS is shown with an outlined arrow, a Ki-67^hi^ RS with a filled arrow. Scale bars = 20 µm. * = p < 0.05, ** = p < 0.01, *** = p < 0.001, ns = non statistically significant.

### Circulating RS are common and clinically relevant in MDS-RS

Reagent-free magnetic isolation was tested in peripheral blood (PB) samples to detect circulating siderocytes. Surprisingly, M+FACS assessment of PB HD cells identified a substantial number of RS (**Fig. 3A**). No previous reports have described appreciable numbers of circulating RS in MDS-RS PB.^4^ However, peripheral RS were a ubiquitous observation correlating well in abundance with BM RS isolates (**Fig. 3B**). Where matched clinical data were available, PB and BM RS isolates were positively correlated with diagnostic smear RS (**Fig. 3C**) and serum erythropoietin (**Fig. 3D**), and negatively correlated with hemoglobin (**Fig. 3E**). Interestingly, the identification of PB RS in FC also detected increased DNA content RS in erythropoiesis-stimulating agent (ESA)-treated patients (**Fig. 3F**). This was morphologically validated in several cases (**Fig. 3G/H**), identifying a positive effect of ESAs on RS (increasing survival of dysplastic cells or increasing mitosis).

**Figure 3:**
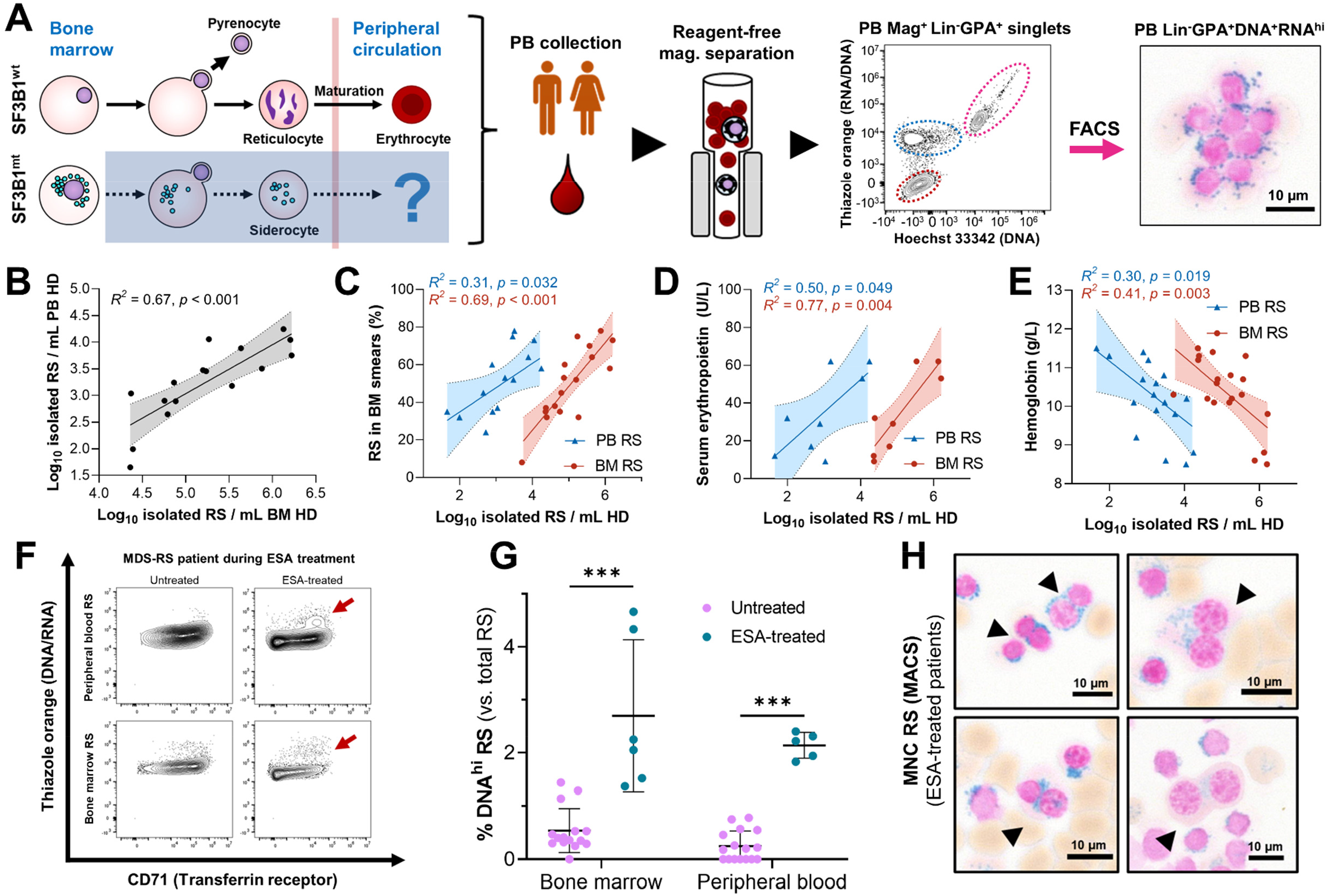
Peripherally circulating RS are common and clinically relevant in MDS-RS. **A)** Isolation steps from the PB HD fraction of MDS-RS patients through reagent-free magnetic separation and representative flow cytometry diagram, where RS are identifiable and validated as present through morphological analysis. **B)** Correlation of RS abundances isolated from matched BM and PB samples (leftmost subpanel, n = 16). **C-E)** Correlation of log_10_-converted isolated RS numbers obtained from anemic (Hb<12.0 g/dL) MDS-RS patients with **C)** BM RS percentages from clinical BM smears (n^PB^ = 15, n^BM^ = 17), **D)** hemoglobin levels (n^PB^ = 18, n^BM^ = 19) and **E)** serum erythropoietin levels (untreated patients only, n^PB^ = 8, n^BM^ = 8). **F)** Flow cytometry example of BM and PB RS with increased DNA content comparing two visits of the same patient to the clinic, before and after ESA treatment. A cell population of increased DNA content is highlighted with dark red arrows. **G)** Mean (SD) frequency of RS with elevated DNA content, separated by EPO treatment status and cell fraction of origin. Arrows indicate binucleate RS identified during morphological analysis of EPO-treated and RS-enriched samples (scale bars = 10 µm). **H)** Morphological visualization of binucleated RS in ESA-treated, RS-enriched samples (scale bars = 10 µm). * = p < 0.05, ** = p < 0.01, *** = p < 0.001, ns = non statistically significant.

### SF3B1 mutations have limited impact on the gene expression of true MDS-RS HSPC

We next investigated *SF3B1*^mt^ erythropoiesis through RNA sequencing, comparing MDS-RS cell populations against healthy individuals at bulk (**Fig. 4A, Table S1**) and single-cell level (**Fig. 4B, Fig. S8-S12, Data S1**). We first focused on CD34^+^ HSPC, as these constitute the compartment of origin in MDS-RS.^6^ As previously reported,^10,20^ differential gene expression (DE) analysis comparing MACS-enriched CD34^+^ cells between MDS-RS and healthy individuals identified overexpression (OE) of erythroid genes and underexpression (UE) of known *SF3B1*^mt^ targets (**Fig. 4C**, left panel; **Data S2**). However, no increased erythroid transcripts were observed in transcriptomically-identified HSPCs (**Fig. 4C**, right panel). AUCell analysis^21^ was then conducted to assess a potential proerythroid signature (top 100 EB development genes),^22^ and similarly demonstrated no difference between MDS-RS and NBM HSPC clusters (**Fig. 4D**).

**Figure 4:**
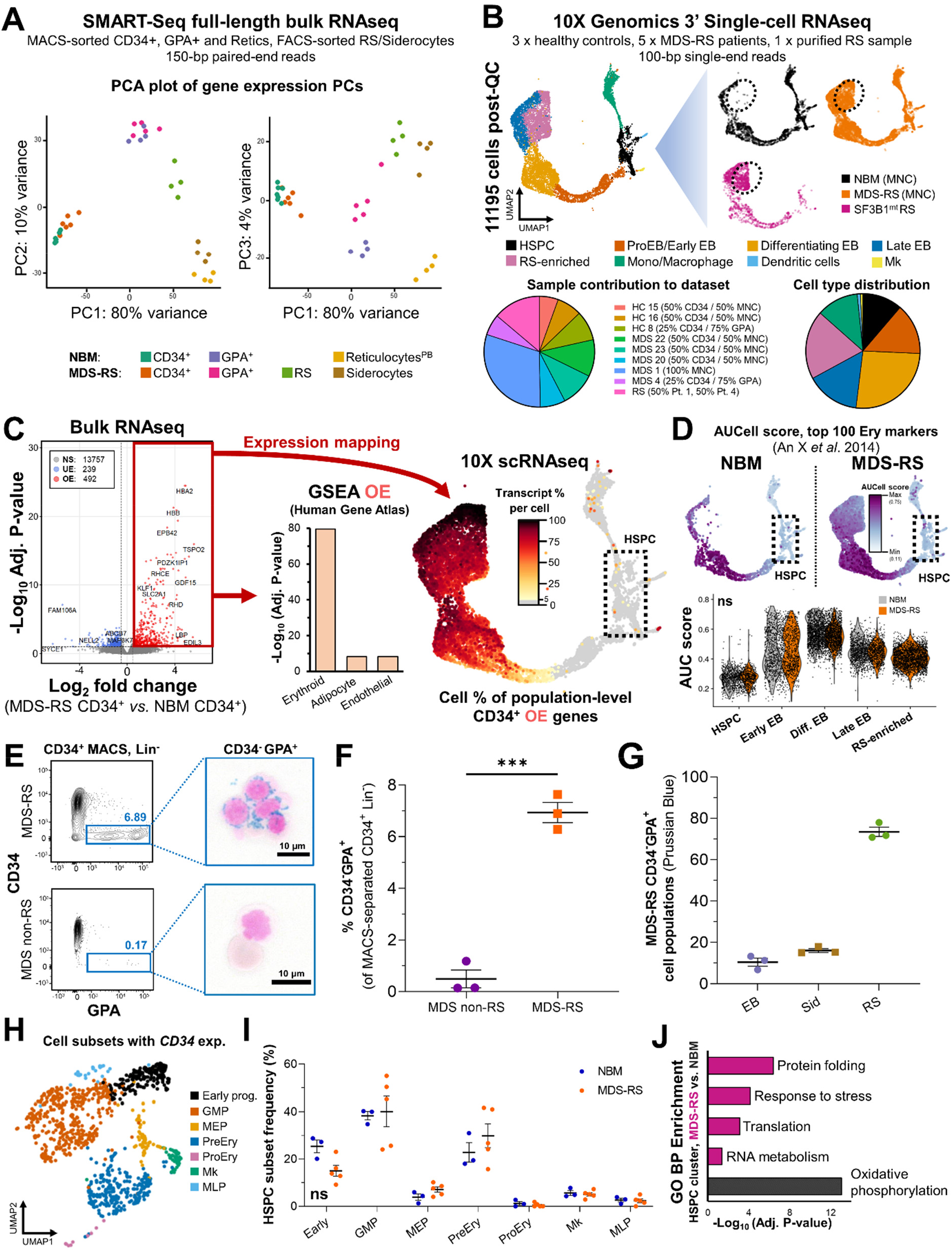
*SF3B1* mutations have limited impact on the gene expression of true MDS-RS HSPC. **A)** Principal component analysis (PCA) plots of a full-length bulk RNAseq experiment encompassing sorted cell populations from NBM donors (n^CD34^ = 7, n^GPA^ = 4, n^Ret^ = 4) and *SF3B1*^mt^ MDS-RS patients (n^CD34^ = 6, n^GPA^ = 5, n^RS^ = 4, n^Sid^ = 4). Sample distribution along PC 1 is visualized against PC 2/PC 3. **B)** Global overview of 2 integrated 10X Genomics scRNAseq experiments encompassing sorted cell populations from NBM donors and *SF3B1*^mt^ MDS-RS patients (n^NBM^ = 3, n^MDS-RS^ = 5, n^RS^ = 1). A UMAP-based bidimensional projection is displayed and separated per sample group, where each cell is visualized as one point. The dotted circle indicates the RS-enriched cell subset which is absent in NBM. Cell types were annotated according to gene expression signatures per cluster set. Sample and cell type composition in the total dataset are shown below the UMAP plots. **C)** Volcano plot (**left**) displaying differentially expressed genes (DEG) in CD34^+^ MACS-enriched BM cells comparing *SF3B1*^mt^ MDS-RS vs. NBM. Cut-offs for significance were Log_2_ FC > 0.5, adjusted P-value < 0.01. Genes were overexpressed (OE, red), underexpressed (UE, blue) or not significantly different (NS, grey). Gene set enrichment analysis (GSEA) of OE genes (middle) was performed with the Enrichr Human Gene Atlas. The **right** UMAP heatmap displays expression of bulk OE genes in scRNAseq. The dotted rectangle highlights the HSPC transcriptomic cluster. **D)** AUCell erythroid score (based on erythroid markers from An et al. 2014^22^) mapped in the UMAP overlays and separated by mutational background. The erythroid score is similarly displayed in violin plots (NBM in grey, *SF3B1*^mt^ in orange) and grouped per cell population (excluding cell subsets unrelated to erythroid development, e.g. macrophages). **E)** Representative CD34 and GPA FACS plots from CD34^+^ MACS-separated BM MNCs isolated from an *SF3B1*^mt^ MDS-RS patient and from a non-*SF3B1*^mt^ non-RS MDS patient. Lin^-^CD34^-^GPA^+^ cells are gated in blue and connected to representative micrographs. Scale bars = 10 µm. **F)** Mean (± SEM) percentage of Lin^-^CD34^-^GPA^+^ cells in CD34^+^-enriched cells (n = 3 per group). **G)** Mean (± SEM) cell frequencies based on morphological analysis of Lin^-^CD34^-^GPA^+^ in MACS-purified CD34^+^ MDS-RS samples. **H)** UMAP projection of *CD34* RNA-positive HSPCs in the scRNAseq dataset. **I)** Mean (± SEM) frequencies of transcriptomically identifiable HSPC subsets as set out in panel **H** and compared between NBM and MDS-RS samples. **J)** Gene set enrichment analysis results for Gene Ontology Biological Process (GO BP) enrichment of DEG identified in the HSPC cluster between MDS-RS and NBM cells (n^NBM^ = 432, mean 144 cells/donor; n^MDS-RS^ = 510, mean 102 cells/donor). * = p < 0.05, ** = p < 0.01, *** = p < 0.001, ns = non statistically significant.

In light of the efficiency of RS isolation with reagent-free MACS, we investigated whether antibody-mediated MACS enrichment could induce sample contamination. Indeed, CD34^+^-enriched MDS-RS cells displayed a Lin^-^CD34^-^GPA^+^ population averaging ∼90% RS/siderocytes which was absent in MDS samples without RS (**Fig. 4E-G**), highlighting important limitations in interpreting MACS-enriched cell data from MDS-RS.^8-11^ To circumvent these, we evaluated the molecular signature of *CD34* transcript-positive HSPC in scRNAseq (**Fig. 4H, Fig. S13**, **Data S3**). Transcriptomically-defined HSPC subset frequencies were unchanged (**Fig. 4I**), matching previous data of phenotypical HSPC in MDS-RS^6^. Interestingly, few DE genes were detected in CD34^+^ HSPCs despite these maintaining RNA mis-splicing, and the only enrichment identified was a protein/ER stress-associated response (**Fig. 4J**). Given these limited HSPC results, we next focused our approach on specifically investigating erythroid biology.

### RS actively modulate their gene expression to survive extensive oxidative/splicing stress

RS were used as the primary point of comparison against NBM EB due to being phenotypically and clonally engaged. Bulk RNAseq identified extensive underexpression of cell cycle-related genes, matching our initial characterization (**Fig. 5A**, **Data S3**). Most importantly, overexpressed genes in bulk RNAseq mapped to an MDS-specific scRNAseq cluster which was over-represented in RS and absent in healthy donors (**Fig. 5B**). This cluster presented a more aberrant transcriptomic phenotype in purified HD-RS versus MNC-RS (**Fig. 5C**). However, and beyond this MDS-specific cluster, RS clearly encompassed all stages of erythroid differentiation.

**Figure 5:**
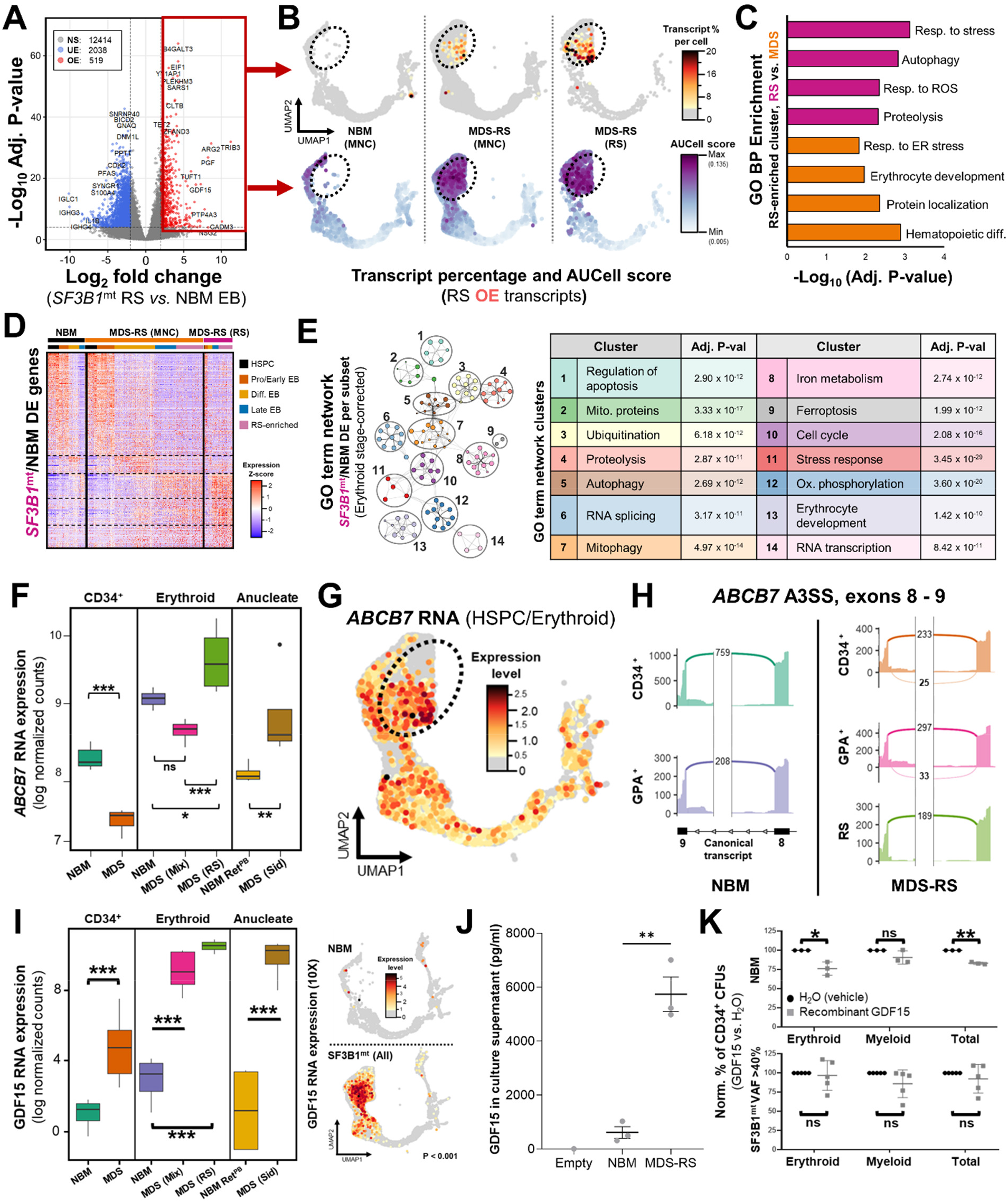
*SF3B1^mt^* RS activate a pro-survival transcriptomic program against oxidative and RNA splicing stress. **A)** Volcano plot displaying differentially expressed genes (DEG) in bulk data comparing M+FACS-purified *SF3B1*^mt^ RS against MACS-purified NBM GPA^+^ erythroblasts with an absolute Log_2_ FC cut-off of 2 and an adjusted P-value cut-off of 10^-4^. **B)** UMAP overlays of RS OE transcript percentages per cell (top row) and AUCell scores of RS identity (bottom row), separated by sample type. Transcript percentages are mapped with an initial baseline cut-off of 4% of total transcripts. AUCell scores were based on the RS OE gene set. **C)** GO biological process (BP) enrichment analysis of DEG identified in the RS-enriched cluster comparing HD fraction-derived RS versus MNC-derived RS (non-specifically present among MDS-RS MNCs and with presumably decreased iron load). **D)** Heatmap of all differentially expressed genes between RS from *SF3B1*^mt^ MDS-RS patients vs. NBM, subclustered by cell subset. The upper bar above the heatmap identifies each sample type, and cells are further clustered according to cell type, identified by the lower bar. Dashed lines highlight cell type separation. **E)** Metascape GO term network generated from all DEG identified through comparison of RS and NBM samples at each transcriptomically-identified differentiation stage cluster (HSPC, ProEB + Early EB, Differentiating EB and LateEB + RS-enriched), correcting for differentiation stage skewing. GO sub-terms (small circles) are organized and clustered by major functional terms (numbered black circles). Clusters are annotated in the table to the right, including adjusted P-values from Metascape analysis. **F)** *ABCB7* RNA expression in bulk RNAseq, displayed in log normalized counts from all assayed cell populations. **G)** Gene expression values of *ABCB7* overlaid in the UMAP projection, with grey cells displaying no detectable expression and a gradient from light yellow to dark red indicating the level of gene expression per cell. **H)** Sashimi plots for canonical (normal font) and mis-spliced (bold) read counts of the *ABCB7* alternative 3’ splice site associated with targeting by NMD. **I)** *GDF15* expression based on RNA sequencing of purified populations (**left**, quantified in log normalized counts) and single cells (**right**, UMAP overlay). **J)** Mean (± SEM) GDF15 concentration in culture supernatants obtained from 28-day erythroid culture of BM MNCs (n^NBM^ = 3, n^MDS-RS^ = 3), as determined by ELISA. 1 empty scaffold was kept in the same media and culture conditions to evaluate GDF15 levels in base media. **K)** Mean (± SD) erythroid and myeloid colony formation from MACS-enriched CD34 cells (n^NBM^ = 3, n^MDS-RS^ = 5), normalized to untreated numbers. Minimum total colonies counted were 254 among NBM donor conditions and 124 among MDS-RS donor conditions. Cells were treated with either recombinant GDF15 peptide at a concentration of 100 ng/ml (grey squares) or with an equal volume of water (vehicle, black circles). * = p < 0.05, ** = p < 0.01, *** = p < 0.001, ns = non statistically significant.

DE analysis was then performed between RS/NBM within each scRNAseq transcriptomic cluster to define differentiation-independent alterations (**Fig. 5D, Data S4**). This approach identified a major enrichment of stress response pathways (**Fig. 5E**), particularly related to autophagy, proteasomal processing and mitophagy. Metabolic changes were also evident, particularly the significant enrichment of several antioxidant pathways (glutamine/ROS response); together, these indicate an active transcriptional state in RS to maintain cell homeostasis against overactive mis-splicing and oxidative stress.

Erythroid-specific genes were also heavily affected in this comparison, with RS displaying loss of expression of hemoglobin chains *HBA2* and *HBD*, a hallmark for iron deficiency anemia^23^ (**Fig. S14**), and decreased *HBB* expression in cells with higher RS-associated transcripts (**Fig. S15**). Conversely, erythroid gene expression was higher in peripherally circulating RS, suggesting that this population is either more differentiated or less phenotypically affected (**Fig. S16/S17**).

Focusing on bulk expression of *ABCB7*, the major *SF3B1*^mt^-induced NMD-targeted event in *SF3B1*^mt^ MDS-RS,^10,20^ we found that its abundance was evidently decreased in HSPCs (matching previous reports) but significantly increased in RS/siderocytes (**Fig. 5F**). We cross-validated this observation in scRNAseq RS (**Fig. 5G, Fig. S18**) and identified lower levels of the cryptic *ABCB7* splice site in RS as compared to CD34^+^ and GPA^+^ cells (**Fig. 5H**).

Both RNAseq datasets identified overexpression of *GDF15* (Growth Differentiation Factor 15) as a major occurrence (**Fig. 5I**). *GDF15* is a stromal/erythroid stress factor previously shown to be increased in MDS-RS patient sera^24^ and reported to suppress HSPC growth in culture.^25^ When *in vitro* culture was used to induce stress erythropoiesis, cultured *SF3B1*^mt^ cells still secreted more than 9X GDF15 than normal cells, highlighting *GDF15* overexpression is disease-related (**Fig. 5J**). A colony-forming unit (CFU) experiment to evaluate GDF15 effects on HSPC suppression displayed variable results in MDS-RS samples but identified a consistent and significant decrease in NBM erythroid/total CFU forming ability (**Fig. 5K**), linking GDF15 secretion by RS to an effect on upstream HSPC biology.

### SF3B1^mt^ mis-splicing is context-specific and significantly increased in RS

Given the surprising patterns of *ABCB7* expression/splicing in RS (**Fig. 5F-H**), we next focused on a broader analysis of alternative splicing (AS) in the bulk RNAseq data comparing healthy individuals and *SF3B1*^mt^ MDS-RS (HSPC, RS vs. EB [nucleated erythroid (N)] and siderocytes vs. reticulocytes [anucleate erythroid (A)]. RS presented substantially more differential AS events; 63 genes were AS in both HSPC/RS, and only 7 genes were AS in all 3 groups (**Fig. 6A**).

**Figure 6:**
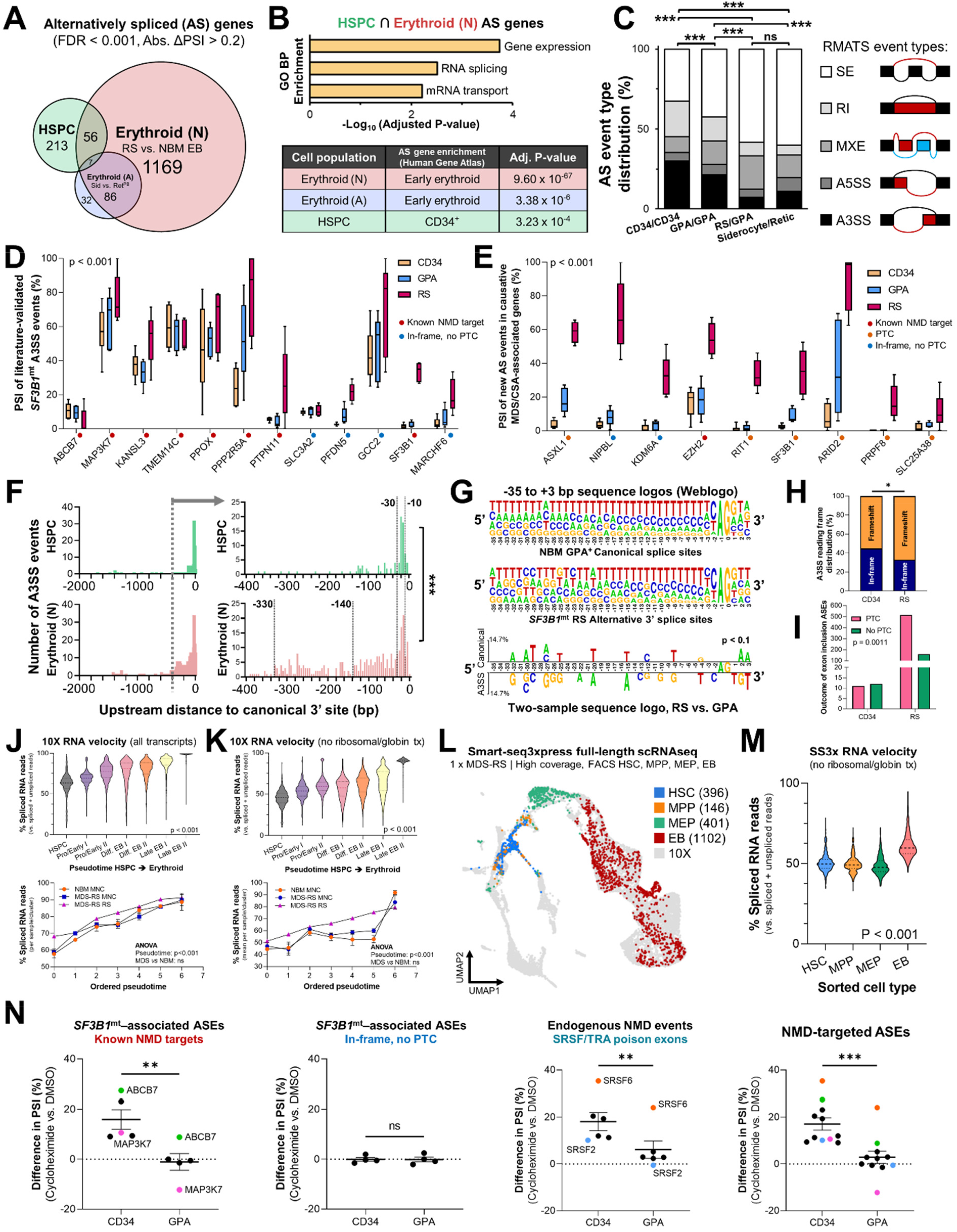
Distinct RNA dynamics in erythroid differentiation intensify *SF3B1*^mt^ mis-splicing. **A)** Proportional Venn diagram of genes undergoing statistically significant alternative splicing (AS; false discovery rate [FDR] < 0.001, absolute difference in percentage spliced in levels [Abs. ΔPSI] > 0.2) in the HSPC (green, MDS-RS CD34^+^ vs. NBM CD34^+^), nucleated erythroid (red, MDS-RS RS vs. NBM EB, *Erythroid (N)*) and anucleate erythroid (blue, MDS-RS Siderocytes vs. NBM Ret^PB^, *Erythroid (A)*) populations. **B)** GO BP enrichment analysis results comprising genes mis-spliced in both the HSPC and Erythroid (N) populations are shown in the upper bar chart. The Human Gene Atlas enrichment result with the lowest adjusted P-value is shown in the lower table. **C)** Frequency of AS events split by rMATS category in each sample group comparison (SE = skipped exon, RI = retained intron, MXE = mutual exon exclusion, A5SS = alternative 5’ splice site, A3SS = alternative 3’ splice site). Statistical comparisons of A3SS+RI frequencies were performed with Fisher’s exact test. **D)** Box plots of percent spliced-in (PSI) values of literature-validated *SF3B1*^mt^-induced A3SS events in MDS-RS samples, separated by sample type (CD34, GPA, RS). Known targeting by NMD is indicated with a red circle, in-frame events without a premature termination codon (PTC) are indicated by a blue circle. **E)** Box plots of PSI values in newly-identified ASEs affecting known MDS and congenital sideroblastic anemia (CSA) causative genes. Known targeting by NMD is indicated with a red circle, PTC detection with unverified NMD is indicated with an orange circle, in-frame events without a PTC are indicated by a blue circle. **F)** Distribution of base pair distances from cryptic A3SS sites to canonical splice sites (horizontal axis) in HSPC and Erythroid (N). Further detail is provided in -400 bp to 0 bp for increased contrast. Lines at -30 bp and -10 bp demarcate the interval associated with *SF3B1* mis-splicing.^30^ Additional lines at -140 bp and -330 bp demarcate additional erythroid intervals of interest. **G)** Sequence logos of canonical and A3SS sequences encompassing the 3’ splice site (starting at -35 bp upstream of the AG motif) and statistical comparison through a two-sample logo^47^. **H)** Frequency of A3SS events per rMATS cell type comparison where the splice site shift remains in-frame (blue) or induces a frameshift event (orange). **I)** Frequency of exon insertion events per rMATS cell type comparison where the splice site shift incorporates a new PTC (pink) or remains in-frame with no PTC induction (green). **J)** RNA velocity analysis of transcriptomically-identified HSPC and EB subsets in 10X scRNAseq, visualizing the percentage of spliced transcripts along pseudotime in the total cell populations (violin plots) or separated by sample group (scatter plots). The left column quantifies all transcripts, whereas the right column excludes ribosomal and globin transcripts. **K)** UMAP overlay of FACS-purified HSPC subsets and GPA+ EB from 1 *SF3B1*^mt^ MDS-RS patient after Smart-seq3xpress (SS3x), visualizing true vs. predicted cell type identity. **L)** RNA velocity analysis of spliced RNA read percentages in the FACS-sorted SS3x experiment, analyzed independently of the 10X dataset. This graph excludes ribosomal and globin transcripts. **M)** Mean (± SEM) differences in PSI after 3 h cycloheximide treatment (70 µg/mL) versus DMSO (1:1000, vehicle) in MDS-RS CD34 and GPA cells. *SF3B1*^mt^-associated NMD-targeted ASEs with sufficient coverage are shown at far left, *SF3B1*^mt^-associated in-frame ASEs at middle-left, and endogenous NMD-targeted transcripts^48^ at middle-right. The far-right plot visualizes all ASEs. * = p < 0.05, ** = p < 0.01, *** = p < 0.001, ns = non statistically significant.

Shared AS events between HSPC/RS were enriched in RNA splicing-associated genes (**Fig. 6B**), in line with previous *SF3B1*^mt^ studies.^10,26^ However, genes that were specifically AS in each population significantly enriched for cell type identity, reflecting distinct transcriptomic programs. AS event types also progressively shifted toward predominantly exon skipping (SE) events during erythroid differentiation, compared to the initial skew of alternative 3’-splicing (A3SS) and intron retention (RI) events in HSPCs (**Fig. 6C, Fig. S19**).

Focusing on literature-validated *SF3B1*^mt^ AS events^10,27^, we found that decreased mis-splicing of *ABCB7* was a singular case in RS; in fact, every other AS event displayed a constant or increased median level of the cryptic transcript in RS/EB as compared to HSPC, including known NMD-targeted AS (**Fig. 6D**). New AS sites affecting several MDS-causative genes^28^ and *SLC25A38* (causative in congenital sideroblastic anemia^29^) were detected in HSPC at very low levels and, to our knowledge, previously unreported. However, mis-splicing in these genes increased in EB to become the predominant transcript form in RS, with many of these inducing frameshifts and/or inclusion of premature termination codons (PTC) (**Fig. 6E**).

Functional analysis of cryptic A3SS site locations in HSPCs confirmed their enrichment between - 30 and -10 bp upstream of the canonical 3’ site (**Fig. 6F**), as previously reported.^30^ Conversely, RS displayed events in the same interval but had a much larger distance distribution, indicating reduced fidelity for splice site recognition. This is functionally supported by differentially altered RS nucleotide frequencies upstream of and surrounding the cryptic 3’ splice site (**Fig. 6G**), as well as a much larger proportion of frameshift/PTC events caused by A3SS/SE events (**Fig. 6H/I**).

### Modified RNA dynamics in erythropoiesis justify the magnified mis-splicing profile of RS

We next hypothesized that the highly aberrant AS profile in RS could be caused by altered splicing/NMD dynamics in erythropoiesis. RNA velocity analysis^31^ of spliced transcript percentages in the 10X clusters identified increased splicing rates even in early EB (**Fig. 6J**). We were concerned that the high erythroid transcript burden of ribosomal/globin genes could skew these data; however, this trend remained after discarding these transcripts from analysis (**Fig. 6K**). Importantly, NBM and MDS samples behaved identically, highlighting the role of differentiation and not disease in this process. As velocity analysis of the 10X samples was based on transcriptomically-identified clusters, we then confirmed the same splicing dynamics in FACS-purified HSPC/EB populations through Smart-seq3xpress (**Fig. 6L/M, Fig. S20-S22**).

Cycloheximide treatment of HSPC and EB was then pursued to evaluate NMD dynamics as originally reported in HSPC by Shiozawa et al.^9^ Overall, NMD-targeted cryptic transcripts were increased in frequency by cycloheximide treatment in HSPC but not in EB (**Fig. 6N**), through which we determine decreased NMD activity during erythroid differentiation. This is consistent with the significantly higher proportions of NMD-targeted AS events in RS (**Fig. 6D**). Thus, taken together, we conclude that erythroid differentiation magnifies the downstream effects of *SF3B1*^mt^.

### Integrative proteomic analysis of SF3B1^mt^ RS identifies disruption of proapoptotic genes

Semi-quantitative proteomics of healthy and MDS-RS erythroblasts/RS were performed and integrated with RNAseq to functionally assess the outcome of *SF3B1*^mt^ expression/splicing (**Fig. 7A**, **Data S5**). This analysis identified four major signatures of differential RNA/protein expression which validated our previous scRNAseq results (**Table S2, Data S5**). The same pro-survival pathways from scRNAseq were validated as overexpressed, together with a highly significant decrease in cell cycle-associated proteins; however, the proteomic dataset also identified a specific and significant decrease in NMD target-degrading proteins (**Fig. S23**). Importantly, GDF15 was significantly overexpressed in both EB and RS, further supporting previous findings (**Fig. 7B**).

**Figure 7:**
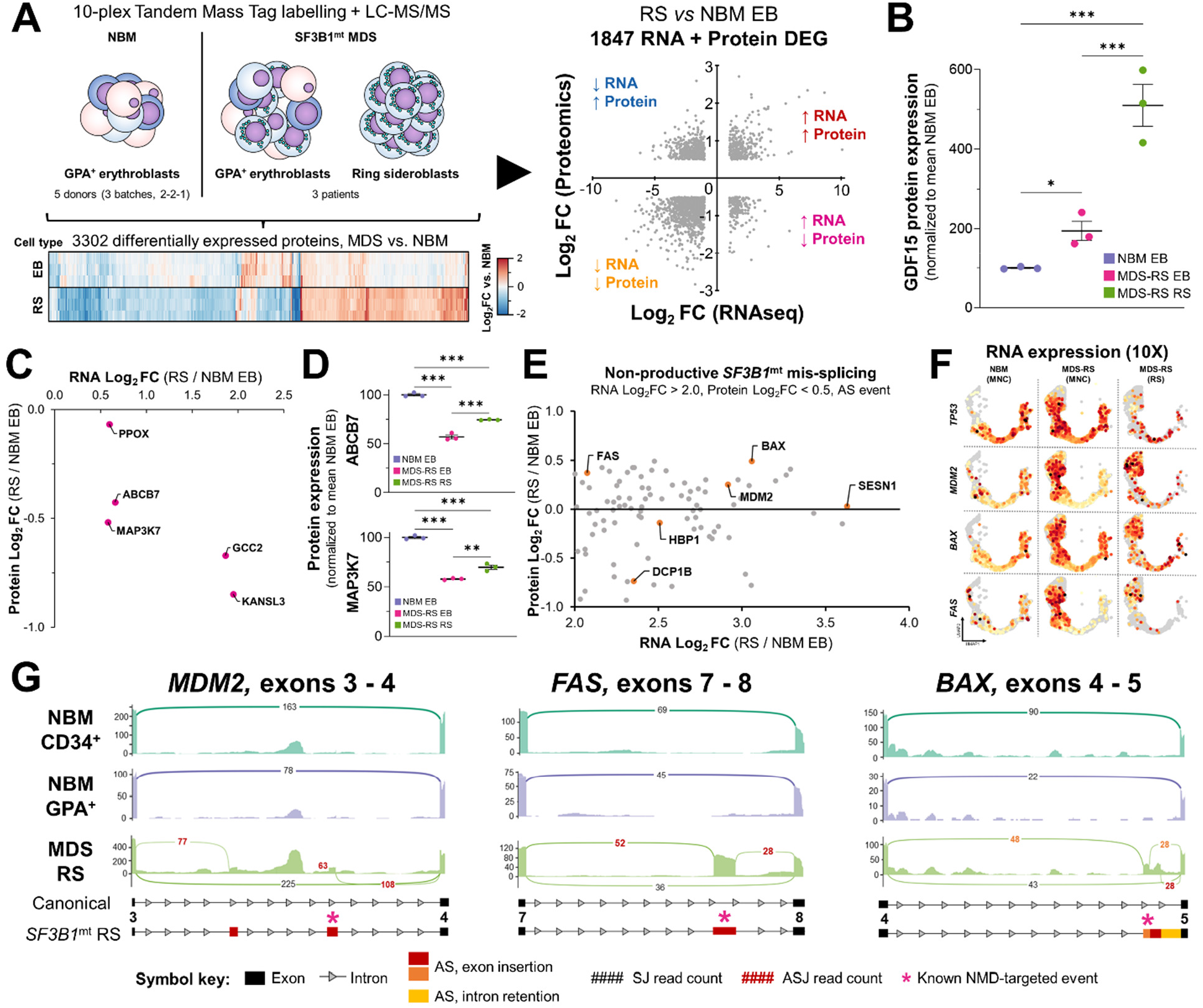
Proteomic analysis of *SF3B1*^mt^ RS defines mis-splicing errors affecting proapoptotic genes. **A)** Design of a combined transcriptomic and proteomic analysis of *SF3B1*^mt^ RS. EB samples from 5 NBM donors (separated into 3 biologically distinct batches) and paired EB + RS samples from 3 MDS-RS patients were subjected to semi-quantitative proteomics. DEG are compared against differentially expressed proteins (DEP) to obtain four major signatures of differential expression, highlighted in each quadrant and expanded on in **Table S2**. **B)** Mean (± SEM) protein expression level of *GDF15*, normalized to mean NBM expression. **C)** Scatter plot of literature-validated SF3B1^mt^ mis-spliced genes where increased RNA expression is accompanied by decreased protein expression in RS. **D)** Mean (± SEM) protein expression levels of *ABCB7* and *MAP3K7*, normalized to mean NBM expression. **E)** Scatter plot of genes detected to be both significantly AS and OE in RNAseq data and without significant increase in protein expression, predicted as undergoing non-productive mis-splicing events in RS. *TP53* pathway genes were detected by enrichment analyses and are highlighted in orange with gene symbols included. **F)** Gene expression values of *TP53*, *MDM2*, *BAX* and *FAS* are overlaid in the HSPC/erythroid UMAP projection and separated by sample type, with grey cells displaying no detectable expression and a gradient from light yellow to dark red indicating expression per cell. **G)** Sashimi plots comprising mis-spliced transcript regions of MDM2, FAS and BAX. Canonical splice junction counts (SJ) are noted in black, and alternative SJ counts are noted in red. A full legend is provided below the graph. The asterisks indicate sites corresponding to transcripts known to be targeted by NMD. * = p < 0.05, ** = p < 0.01, *** = p < 0.001, ns = non statistically significant.

Cases of RNA/protein uncoupling (increased RNA expression and decreased protein expression) were particularly interesting as candidate targets of dysfunctional AS events. Indeed, this gene subset (RNA↑Protein↓) encompassed several known *SF3B1*^mt^-affected genes*^10,30^* (**Fig. 7C**). Investigating *ABCB7* and *MAP3K7* protein levels demonstrated significant underexpression in MDS-RS EB and RS vs. healthy EB. However, both proteins were significantly increased in RS vs. MDS-RS EB (**Fig. 7D**), indicating RS compensate mis-splicing via RNA overexpression.

We thus expanded our search of non-productive mis-splicing to include further cases of RNA/protein uncoupling by prioritizing significantly AS genes with high RNA overexpression and low correlation to protein abundance. Interestingly, this resulted in specific enrichment of the *TP53* pathway (**Fig. 7E**). Due to their key role in apoptosis, *MDM2*, *BAX and FAS* were further investigated and validated as overexpressed in scRNAseq (**Fig. 7F**). Diverse NMD-associated events barring functional protein production were identified in both RNAseq datasets as exclusive to RS (rMATS FDR < 10^-5^, **Fig. 7G, Fig. S24**). We thus conclude that apoptotic regulation is abnormal in RS due to RNA mis-splicing, and for the first time connect *SF3B1* mutations to dysfunction of the TP53 pathway.

## Discussion

Whilst *in vitro* and *in vivo* studies of *SF3B1^mt^*erythropoiesis have proven successful in modelling HSPC pathobiology and recreating RS generation, the molecular mechanisms driving the growth and survival of *SF3B1*^mt^ RS in the MDS-RS BM and the competitive advantage of *SF3B1^mt^* cells have remained elusive.^6,15,16^ Through an integrative multiomics approach, we investigated *SF3B1*^mt^ MDS-RS erythropoiesis at a cell and molecular level to address these open questions.

We established a novel method for viable RS and siderocyte isolation from patient BM samples, enabling exploration of their disrupted erythroid development and rendering us the unique possibility to explore the entire process of *SF3B1*^mt^ erythroid development from HSC to RS. This method was also validated in PB samples, identifying circulating RS as a relevant clinical observation correlating in abundance with BM RS burden and hemoglobin/serum EPO levels.

An important outcome associated with reagent-free magnetic isolation of RS was the finding that CD34^+^ MACS enrichment (a core experimental approach for HSPC isolation^8-11,20^) indirectly isolates a significant number of RS and siderocytes due to their high iron content, creating an artificial erythroid subpopulation which was absent in matched MDS samples without RS. This subpopulation fundamentally underlies the bulk transcriptomic erythroid signature conventionally associated with MDS-RS HSPCs, which we also observed in our bulk RNAseq data and yet failed to verify in a detailed scRNAseq analysis including enriched HSPCs. Indeed, despite showing a clear functional impairment in long-term culture-initiating cell and CFU assays,^6^ true MDS-RS HSPCs displayed few alterations in gene expression and no detectable differences in commitment, subset frequencies or erythroid identity. Given this contradiction, we speculate that investigating RNA splicing and protein production at HSPC subset resolution will be instrumental towards understanding the functional consequences of *SF3B1*^mt^ and downstream RNA mis-splicing on early hematopoietic development.

We demonstrated that RS constitute a living, differentiating and molecularly active population, with decreased cell cycle progression as the major sign of dysfunction. RS were found to engage diverse homeostatic mechanisms to survive oxidative and RNA splicing stress, including the surprising protein rescue of known mis-spliced and NMD-targeted genes *ABCB7* and *MAP3K7*.^10^ We explored the molecular mechanisms underlying these changes and define an intensified *SF3B1*^mt^ mis-splicing panorama in RS. We speculate that modified RNA splicing and NMD dynamics in erythroid cells combine to switch “quality control” from RNA to protein. Indeed, RS sensitivity to proteasome inhibition by bortezomib has been demonstrated in a clinical trial.^32^ We directly associate *SF3B1* mutations with disruption of proapoptotic genes in the *TP53* pathway, a finding which may potentiate the malignant *SF3B1* risk profile in *TP53*-mutant settings, e.g. MDS^5q-^ or AML,^33,34^ and increases the need to evaluate how complex genetic profiles potentiate new splicing interactions.

Chronic myeloproliferative neoplasms develop over decades, and we recently reported preliminary data indicating this is also the case in MDS-RS.^35,36^ *SF3B1*^mt^ cell effects on their surroundings may thus comprise a major factor in driving clinical disease. We identified *SF3B1*^mt^ RS as a key source of GDF15 and confirmed a detrimental impact of this cytokine on wildtype HSPC biology. It is thus tantalizing to speculate that RS could function as disease-augmenting “foot soldiers”, similarly to Reed-Sternberg cells in Hodgkin lymphoma,^37^ and further exploration of how *SF3B1*^mt^ cells affect their surrounding microenvironment and wildtype cells will be essential.

In conclusion, our characterization of *SF3B1*^mt^ erythropoiesis constitutes a unique platform for the study of MDS-RS, providing novel insights into the unexpectedly active biology of the “dead-end” RS and enabling further investigation of disease pathogenesis and treatment avenues.

## Methods

### Study design, sample collection and ethical approval

Bone marrow (BM) and/or peripheral blood (PB) samples were collected from 36 MDS-RS and 3 MDS non-RS patients evaluated at Karolinska University Hospital, Huddinge, Sweden. Diagnostic procedures were performed according to the European LeukemiaNet recommendation and WHO classification for myeloid neoplasms.^38,39^ As the specific purpose was to dissect the pathobiology of *SF3B1*^mt^ MDS-RS, all MDS-RS patients belonged to the *SF3B1*^α^ category.^7^ RS presence was quantified according to standard clinical practice. Additional samples were collected from a total of 40 healthy normal bone marrow (NBM) donors for control purposes. A deidentified donor and experiment index including clinical and mutational status^6^ is provided in **Data S6**. All source material was provided with written informed consent for research use, given in accordance with the Declaration of Helsinki, and the study was approved by the Ethics Research Committee at Karolinska Institutet (2010/427-31/3, 2017/1090-31/4).

### BM/PB sample processing and density gradient separation

Samples were separated by Lymphoprep™ (STEMCELL Technologies) density gradient centrifugation at 400g for 30 min, room temperature (RT), to derive a mononuclear cell layer (MNC) and an erythrocyte-rich high-density layer (HD). MNC were cryopreserved in 50% RPMI 1640 Glutamax (ThermoFisher), 40% inactivated fetal bovine serum (FBS, ThermoFisher) and 10% Dimethyl Sulfoxide (DMSO) (Sigma-Aldrich). Cryopreserved MNC were thawed in RPMI 1640 Glutamax + 20% FBS + 100 U/mL DNase I (Sigma-Aldrich). HD cells were washed with PBSAG (phosphate buffer saline [PBS, Sigma-Aldrich] + 1 mg/ml bovine serum albumin [BSA, Sigma-Aldrich] + 2 mg/ml glucose [Sigma-Aldrich]) and stored at 4°C for a maximum of 1 week.

### RS isolation and antibody-mediated magnetic-activated cell sorting (MACS)

For RS isolation, packed HD cells were diluted 1:10 with autoMACS® Running Buffer (MACS buffer, Miltenyi Biotec) at 4°C and distributed at 5 mL per LS column (Miltenyi Biotec), or 5×10^6^ BM MNC were thawed, resuspended in 5 mL MACS buffer, and distributed in one LS column. LS columns were washed with 20 mL MACS buffer and eluted with 5 mL MACS buffer. For antibody-mediated separation, the manufacturer’s protocol (Miltenyi Biotec) was followed for separation using CD34, CD235a/GPA or CD71 MicroBeads.

### Flow cytometry analysis and fluorescence-activated cell sorting (FACS)

Cells were analyzed on a CytoFlex S (Beckman Coulter) or analyzed and sorted on a FACS ARIA II Fusion (Becton Dickinson) at the MedH FACS facility of Karolinska Institutet. All steps were performed in FACS buffer (PBS + 2% FBS + 1 mM EDTA) kept at 4°C. All experiments included fluorescent-minus-one (FMO) and single-stained controls. Antibodies/fluorescent reagents utilized are listed in **Table S3**. Data were analyzed using FlowJo v. 10.7.2 (Becton Dickinson).

### Morphological evaluation and microscopy analysis

Cells were cytospun, fixed in methanol for 15 min at RT, air-dried and submitted to Karolinska University Hospital for iron staining. Brightfield micrographs were acquired using a Pannoramic MIDI II slide scanner (3D Histech) at 40x with a Hitachi HV-F22 3CCD SXGA camera (Hitachi Kokusai Electric) using Pannoramic Scanner v. 1.17 (3D Histech). Image analysis was performed using QuPath v. 0.2.0m9^40^ and Fiji v. 2.3.0/1.53f51.^41^ Fixed-cell immunofluorescence is detailed in **Sup. Methods**.

### Droplet digital PCR (ddPCR)

Droplet digital PCR was performed with probes for the detection of *SF3B1* mutations K700E and K666N (Bio-Rad) as previously described.^6^ QuantaSoft analysis software v. 1.7.4 (Bio-Rad) was used to calculate variant allele frequencies (VAF) based on the Poisson distribution. At least one known mutated sample, one wildtype sample and one H_2_O sample were included as controls in every run.

### Bulk RNA sequencing (RNAseq)

CD34^+^ HSPC samples, mixed GPA^+^ erythroblast samples and CD71^+^ PB reticulocyte samples (Ret^PB^) were isolated through MACS. RS and siderocytes were obtained through MACS+FACS. Cells were lysed in RLT (Qiagen) + 40 mM dithiothreitol (Sigma-Aldrich) and RNA extraction was performed with RNeasy Micro Kit (Qiagen) with RNase-free DNase treatment according to the manufacturer’s protocol. RNA integrity numbers (RIN) were estimated using Agilent RNA 6000 Pico Kits (Agilent Technologies, CA, USA). A minimum RIN value of 6.5 was considered adequate. Additional details are provided in **Sup. Methods**. The count matrix and sample metadata are provided in **Data S1**.

### Single-cell RNA sequencing (scRNAseq)

Four experiments were conducted: Two 10X Genomics experiments (5 SF3B1^mt^ MDS-RS patients and 3 NBM donors), one Smart-seq3^42^ experiment (2 MDS-RS donors) and one Smart-seq3xpress^43^ experiment (1 MDS-RS donor). Additional details on sample preparation, sequencing and downstream analysis are provided in **Sup. Methods**. Gene count matrices are under submission for online deposition.

### *In vitro* cell culture and GDF15 measurement

Erythroid culture was initiated by thawing and seeding BM MNCs in polyurethane scaffolds as previously described.^15^ GDF15 levels were measured in both neat (1:1) and diluted (1:10) culture supernatant using a Human GDF15 ELISA kit (LSBio), based on a standard curve of known GDF15 concentrations.

### CD34^+^ colony-forming unit assay

CD34^+^ HSPCs from 3 NBM donors and 5 *SF3B1*^mt^ MDS-RS patients were treated with RPMI plus 100 ng/mL recombinant GDF15 (NBP2-76204-20ug, Novus Biologicals) or 0.5% v/v sterile H_2_O for 1 hour. After preincubation, GDF15 or H_2_O were added to a projected concentration of 100 ng/mL, cells were resuspended in MethoCult™ (H4434; STEMCELL Technologies) and plated in culture dishes. NBM and MDS cells were plated at 4,000 cells/dish and 40,000 cells/dish (due to differential CFU-forming ability^44^), cultured as previously described^6^ and counted using a Leica DM inverted microscope (Leica Microsystems).

### Tandem mass tag (TMT) proteomics

FACS-separated samples were snap frozen in liquid nitrogen. Cell pellets were lysed with 4 % SDS lysis buffer and prepared for mass spectrometry using a modified version of the SP3 protein clean up and digestion protocol.^45^ Additional details are provided in **Sup. Methods** and **Table S4**. Expression matrix (centered against a pool of all samples) files, processed files and GO analyses are provided in **Data S5**.

### Statistical methods

Statistical comparisons were performed through ANOVA, multiple comparison-corrected paired and unpaired T-tests (Benjamini-Hochberg), Fisher’s exact test. Correlation coefficients were calculated and tested for association through Pearson’s product-moment correlation. Statistical methods specific to the analysis of each high-throughput data format are detailed in their respective sections. scRNAseq analysis was performed with RStudio Server v. 1.3.1056 and R v. 3.6.3.^46^ All other statistical analyses were performed with RStudio v. 1.4.1767, R v. 4.0.5, Excel v. 2204 and GraphPad Prism v. 9.4.0.

## Supporting information

Main supplemental file

## Acknowledgments

The authors would like to thank the donors and patients for their willingness to participate in this research. The authors would like to acknowledge support from the MedH Flow Cytometry Core Facility at Karolinska Institutet for providing fundamental assistance with FACS experiments; the facility of Clinical Proteomics Mass Spectrometry at Karolinska University Hospital and Science for Life Laboratory for providing assistance in mass spectrometry and data analysis; the Single-Cell Core Facility (SICOF) at Karolinska Institutet for assistance with Smart-seq3 RS sequencing; and support from the National Genomics Infrastructure in Stockholm funded by Science for Life Laboratory, the Knut and Alice Wallenberg Foundation and the Swedish Research Council, and SNIC/Uppsala Multidisciplinary Center for Advanced Computational Science for assistance with massively parallel sequencing and access to the UPPMAX computational infrastructure.

## Funding

Swedish Cancer Society, 21 0340 (PLM)

Swedish Cancer Society, 19 0200 (EHL)

Knut and Alice Wallenberg Foundation, 2017.0359 (EHL)

Swedish Research Council, 211133 (EHL)

## Authorship contributions

P.L.M. and E.H.L. conceived and designed experiments. P.L.M., T.M-B., I.J.F.H., A-C.B. and I.B. performed cell sorting experiments and associated characterization steps. P.L.M., T.M-B., I.B. and M.C. performed *in vitro* culture experiments. G.T., W.W.K and T.Y. contributed to data analysis. A-C.B., I.B. and G.W. processed and biobanked BM samples. P.L.M., T.M-B. and M.J. performed CFU experiments. M.H-J., C.Z. and R.S. performed SS3xpress scRNAseq, mini-bulk RNAseq, and contributed to design, analysis and interpretation of RNAseq experiments. D.C.G. performed immunofluorescence experiments under supervision of V.L. N.A. and A.J.M. contributed to design and processing of 10X scRNAseq experiments. T.Y., P.S.W. and S.E.W.J. provided age-matched NBM samples. P.L.M., T.M.B., M.D., V.L., P.S.W., S.O., S.E.W.J. and E.H.L. analyzed and interpreted experimental results. P.L.M. analyzed data, created figures, and wrote the manuscript with E.H.L. All authors read, edited, and approved the manuscript.

## Data availability statement

Preprocessed and deidentified RNAseq/proteomic data and downstream results generated in this study are available within the article and its supplementary data files. Raw RNAseq and proteomic data are deposited on a secure Swedish server and have been assigned a DOI (XXX). Data access requests may be submitted to the Science for Life Laboratory Data Centre through the DOI link. Other raw data and analysis code are available from the corresponding author upon request.

