## Supplementary material for "Erythroid differentiation intensifies RNA mis-splicing in *SF3B1*-mutant myelodysplastic syndromes with ring sideroblasts": Main supplemental file

### Main supplemental file - Index

### Supplemental data file description

**Supplemental data 1:** Gene count matrix and experiment metadata for bulk RNAseq

**Supplemental data 2:** DESeq2 differential expression analysis of population-level RNAseq data

**Supplemental data 3:** Seurat clustering and differential expression analysis between RS/NBM and MDS/NBM cell identity clusters in scRNAseq

**Supplemental data 4:** STRING and Metascape enrichment analysis results, respectively separated per cluster and global (corrected by erythroid stage)

**Supplemental data 5:** Semi-quantitative TMT raw data, DE analysis and STRING enrichment analysis comprising NBM EB, MDS-RS EB and purified RS

**Supplemental data 6:** Descriptive statistics, deidentified clinical and experiment map of patients and healthy donors included in the study

### Supplemental methods

**Full-length bulk RNA sequencing and analysis**

RNA sequencing (RNAseq) libraries were prepared from total RNA using SMARTer Stranded Total RNA-Seq Kits v2 - Pico Input Mammalian (Takara Bio, Japan), including enzymatic ribosomal depletion steps. Libraries were sequenced using an Illumina Novaseq 6000 S4 (Illumina, CA, USA) with paired-end 150bp configuration. Reads were pre-processed with TrimGalore v. 0.6.7 using CutAdapt v. 3.5(*1*) and BAM files were generated through via two-pass alignment with STAR v. 2.7.9a(*2*) against the GRCh38.p13 human genome assembly.(*3*) QC details are included in **Sup. Table 2.**

Uniquely mapped read pairs were counted through featureCounts v. 2.0.1(*4*) and downstream gene expression analysis was performed with the use of DESeq2 v. 1.30.1.(*5*) $Log_2$ fold changes ($Log_2FC$) were calculated using DESeq2's maximum likelihood estimate (MLE) method, and associated p-values were calculated using the Wald test and adjusted using the Benjamini-Hochberg (BH) method for multiple testing correction. Gene list enrichment analyses were performed via the STRING database (v. 11.5),(*6*) Enrichr(*7*) Gene Ontology 2021 and Human Gene Atlas modules and the MetaScape server(*8*). Differential splicing analysis was performed between different MDS and NBM cell type groups using rMATS v. 4.1.1,(*9*) with p-values calculated using the likelihood-ratio test (LRT) and adjusted with the BH method. Sashimi plots for visualization were generated using ggsashimi v. 1.1.5.(*10*)

**10X Single-cell RNA sequencing and analysis**

All samples were loaded onto Chromium Single Cell Chips (10x Genomics, CA, USA) at a target capture rate of 10,000 cells per sample. Single cell libraries were prepared using Chromium Next GEM Single Cell 3' Kits v3.1 (10x Genomics) as per the manufacturer's instructions, except 1µl additive ADT primers were added to the initial cDNA PCR amplification buffer and ADT libraries prepared as described in the Total-Seq B protocol (BioLegend) from the initial cDNA SPRI clean up. Libraries were pooled and sequenced on an Illumina NovaSeq 6000 (Illumina). Read pseudoalignment was performed against the GRCh38.p13 human genome assembly (*3*) through kallisto v0.46.1 and bustools v0.40.0(*11*) was used for barcode and UMI counting.

Seurat v3.1.4(*12*) was used to load, process, and analyze the resulting count matrices. For differential gene expression analyses comparing RS and MDS samples against healthy NBM samples, quality control steps were first performed through removal of cells expressing less than 200 distinct genes, total percentage of mitochondrial reads above 15% and a total read count below 2000 counts, as well as through removal of genes expressed in less than 3 cells. All samples were Log normalized and the 2000 most highly variable genes across all samples were identified and used as a basis for integration anchor identification and integration of both datasets. After integration, the data were scaled and Principal Component Analysis (PCA) was performed. The 50 PCs with lowest P-values were selected for UMAP projection(*13*) and clustering (through a shared nearest neighbor [SNN] modularity optimization-based clustering algorithm(*14*) at a resolution parameter of 0.6). The dataset was filtered to include only non-lymphoid hematopoietic cells, and clustering was performed again at a resolution parameter of 2. The generated clusters were then manually annotated by comparison of their gene expression profiles with literature-defined expression patterns and through analysis of the cluster marker genes in Enrichr.(*7*)

Due to the inability to phenotypically identify erythroblast differentiation stages, cells undergoing erythropoiesis were divided into three major stages: early stage erythroid differentiation (proerythroblast [ProEB] and early erythroblast [Early EB]); erythroblasts in active division and differentiation (differentiating erythroblasts [Differentiating EB]); and late-stage terminal differentiation [Late EB]. An additional cluster of late-stage erythroid cells present only in MDS and RS samples was also identified [RS-enriched]. Additional non-lymphoid hematopoietic cell subsets were included in the analysis (monocytes / macrophages [Mono/Macrophage], dendritic cells [DC] and megakaryocytes [Mk]). After cluster identification and annotation, differential expression analysis was performed between sample groups as indicated in each individual comparison through the Seurat-implemented non-parametric Wilcoxon rank sum test, with a $log_2$ fold change threshold of 0.25 and a minimum group-wise percentage of DEG-expressing cells of 10%. The obtained P-values were adjusted with the Bonferroni method and genes with an adjusted P-value below 0.05 were reported as significantly differentially expressed. Gene list enrichment analysis was performed as described above for whole-transcript RNAseq. Due to the relatively low number of CD34-expressing cells in the total dataset, additional integration was performed with literature data obtained from Pellin D. *et al* 2019(*15*) to facilitate cell type mapping. These data were removed after cell type mapping and before differential expression analysis of $CD34^+$ subclusters so as to minimize skewing of the analysis. Z-score heatmaps were generated through use of the DoMultiBarHeatmap function.(*16*)

**Minibulk RNAseq library preparation**

Minibulk RNAseq was performed for assessment of cycloheximide treatment effects in $CD34^+$ and $GPA^+$ cell populations. The library preparation procedure was performed using the Xpress Genomics bulk RNA-seq kit v1, automated on a SP960 liquid handler (MGI Tech). In short, the library preparation procedure denatures RNA samples in presence of oligo-dT primer, which is followed by reverse transcription of RNA with a template-switching procedure and pre-amplification of full-length cDNA for 10 PCR cycles. cDNA was subsequently tagmented using Tn5 (TDE1 Tagment DNA Enzyme; Illumina) and reactions quenched after 10 min at 55 °C by addition of 0.2 % SDS (Sigma-Aldrich). Tagmented DNA was indexed using custom dual-unique Nextera index primers in a 12 cycle PCR reaction. Indexed libraries were cleaned up using SPRI beads in 22% PEG8000 buffer and eluted in 12 µL H2O.

**Smart-seq3xpress library preparation**

Smart-seq3xpress libraries were performed as previously described(*17*) with minor modifications. In brief, cells were sorted into 384-well plates containing 3uL Vapor-Lock (Qiagen) and 0.3uL lysis buffer consisting of 0.125 µM OligodT30VN (5'-Biotin-ACGAGCATCAGCAGCATACGAT$_{30}$VN-3'; IDT) adjusted to reverse transcription (RT), 0.5mM dNTPs/each adjusted to RT volume, 0.1% Triton X-100, 5% PEG8000 adjusted to RT volume, 0.4u RNase Inhibitor (Takara Bio, 40 U/µL). After cell sorting plates were briefly centrifuged before storage at -80C. Before RT, plates were denatured at 72 degrees for 10 min followed by addition of 0.1 µL of RT mix; 25 mM Tris-HCL pH 8.4 (Fischer Scientific), 30mM NaCl (Ambion), 1 mM GTP (Thermo Fisher Scientific), 2.5 mM $MgCl_2$ (Ambion), 8 mM DTT (Thermo Fisher Scientific), 0.25 U/µl RNase Inhibitor (Takara Bio), 0.75 µM Template Switching Oligo (TSO) (5′-Biotin-AGAGACAGATTGCGCAATGNNNNNNNNWWrGrGrG-3′; IDT) and 2 U/µl of Maxima H Minus reverse transcriptase (Thermo Fisher Scientific). Plates were quickly centrifuged after dispensing to ensure merge of lysis and RT volumes. RT was incubated at 42 °C for 90 minutes, followed by ten cycles of 50 °C for 2 minutes and 42 °C for 2 minutes. After RT, 0.6 µL PCR mix was dispensed to each well containing the following; 1× SeqAmp PCR buffer (Takara Bio), 0.025 U/µl of SeqAmp polymerase (Takara Bio) and 0.5 µM Smartseq3 forward and reverse primer. Plates were quickly spun down before being incubated as follows: 1 minute at 95 °C for initial denaturation, 14 cycles of 10 seconds at 98 °C, 30 seconds at 65 °C and 2–6 minutes at 68 °C. Final elongation was performed for 10 minutes at 72 °C.

Smart-seq3 forward primer: 5′-TCGTCGGCAGCGTCAGATGTGTATAAGAGACAGATTGCGCAATG-3′; IDT
Smart-seq3 reverse primer: 5′-ACGAGCATCAGCAGCATACGA-3′; IDT

After PCR, pre-amplified libraries were diluted with 9 µL $H_2O$, before transferring 1 µL of diluted cDNA from each well into a new 384-well plate. Tagmentation was performed by adding 1 µL of tagmentation mix; 1x tagmentation buffer (10 mM Tris pH 7.5, 5 mM $MgCl_2$, 5% DMF), 0.0025 µL Tagmentation DNA Enzyme 1 (TDE1; Illumina DNA sample preparation kit) to the 1 µL of diluted cDNA per well. Plates were incubated for 10 min at 55 °C before the reaction was stopped by the addition of 0.5 µL 0.2% SDS to each well. Index PCR was carried out after the addition of 3.5 µL custom Nextera Index primers (0.5 µM) by dispensing 2 µL of PCR mix; 1× Phusion Buffer (Thermo Fisher Scientific), 0.01 U/µl of Phusion DNA polymerase (Thermo Fisher Scientific), 0.025% Tween-20, 0.2 mM dNTP each. PCR was performed out at 3 minutes at 72 °C; 30 seconds at 95 °C; 14 cycles of (10 seconds at 95 °C; 30 seconds at 55 °C; 30–60 seconds at 72 °C); and 5 minutes at 72 °C. Each indexed library plate was pooled by spinning out gently using a 300-ml robotic reservoir (Nalgene) fitted with a custom 3D-printed scaffold by pulsing the centrifuge to < 200g. The pooled libraries were afterwards purified with in-house 22% PEG beads at a ratio of 1 sample to 0.7 beads.

**Minibulk and Smart-seq3xpress sequencing**

Both minibulk and Smart-seq3xpress libraries were sequenced on a MGI DNBSEQ G400RS platform. Prior to sequencing MGI platform ready circular ssDNA libraries were created using the MGIEasy Universal Library Conversion Kit (MGI). Adapter conversion PCR was carried out on 50ng of final pooled library for 5 cycles, following circularization of 1pmol dsDNA according to manufacturers protocol. DNA nanoballs (DNBs) were created from 80 fmol of circular ssDNA library pools using a custom rolling-circle amplification primer (5′-TCGCCGTATCATTCAAGCAGAAGACG-3′, IDT). DNBs were sequenced PE100 using the custom sequencing primers provided below.

Read 1: 5′-TCGTCGGCAGCGTCAGATGTGTATAAGAGACAG-3′;
MDA: 5′-CGTATGCCGTCTTCTGCTTGAATGATACGGCGAC-3′,
Read 2: 5′-GTCTCGTGGGCTCGGAGATGTGTATAAGAGACAG-3′;
i7 index: 5′-CCGTATCATTCAAGCAGAAGACGGCATACGAGAT-3′;
i5 index: 5′-CTGTCTCTTATACACATCTGACGCTGCCGACGA-3′.

**Minibulk and Smart-seq3xpress data preprocessing**

The zUMIs 2.9.7 pipeline was used to preprocess raw FASTQ files. UMI-containing reads in case of Smart-seq3xpress were identified by the (ATTGCGCAATG) pattern allowing up to two mismatches. Both minibulk and Smart-seq3xpress data was mapped to the human genome (hg38) using STAR version 2.7.3, and readcounts and umicounts were calculated using Gencode v42 annotation.

**RNA velocity analysis**

RNA velocity predictions were performed in 10X and Smart-seq3xpress data through Velocyto(*18*) and further analysed with Scanpy/Scvelo libraries.(*19, 20*) Scanpy-based integration of the two 10X experiments was performed with the Scanorama module.(*21*) The same QC constraints utilized with Seurat were followed with the Scanpy-based analysis. For removal of ribosomal and globin transcripts from the total prediction, gene subsets were excluded from the count matrices before cell QC steps.

**Proteomics sample processing and analysis**

Cell pellets were dissolved in 100 µl Lysis buffer (4% SDS, 50 mM HEPES pH 7.6, 1 mM DTT), heated to 95°C and sonicated. Samples were then prepared for mass spectrometry analysis using a modified version of the SP3 protein clean-up and a digestion protocol,(*22, 23*) where proteins were digested by LysC and trypsin (Pierce sequencing grade modified, Thermo Fisher Scientific).

Each sample was alkylated with 4 mM Chloroacetamide. Sera-Mag SP3 bead mix (20 µL) was transferred into the protein sample together with 100% Acetonitrile to a final concentration of 70 %. The mix was incubated under rotation at room temperature for 18 min. The mix was placed on the magnetic rack and the supernatant was discarded, followed by two washes with 70 % ethanol and one with 100 % acetonitrile. The beads-protein mixture was reconstituted in 50 µL LysC buffer (0.5 M Urea, 50 mM HEPES pH: 7.6 and 1:50 enzyme (LysC) to protein ratio) and incubated overnight. Finally, trypsin was added in 1:50 enzyme to protein

ratio in 50 µL 50 mM HEPES pH 7.6 and incubated overnight. The peptides were eluted from the mixture after placing the mixture on a magnetic rack, followed by peptide concentration measurement (DC Assay, Bio-Rad).
The samples were then pH adjusted using TEAB pH 8.5 (100 mM final conc.), 15 µg of peptides from each sample were labelled with isobaric TMT-tags (TMT10plex reagent) according to the manufacturer's protocol (Thermo Fisher Scientific), and separated by immobilized pH gradient - isoelectric focusing (IPG-IEF) on 3–10 strips as described in Branca R *et al.* 2014.(*24*) The labelling efficiency was determined by LC-MS/MS before pooling of the samples. For the sample clean-up step, a solid phase extraction (SPE strata-X-C, Phenomenex, CA, USA) was performed and purified samples were dried in a SpeedVac. An aliquot of approximately 10 µg was suspended in LC mobile phase A and 1 µg was injected on the LC-MS/MS system. Online LC-MS was performed as described in Branca R *et al.* 2014 using a Dionex UltiMate™ 3000 RSLCnano System coupled to a Q-Exactive-HF mass spectrometer (Thermo Fisher Scientific). Each of the 72 plate wells was dissolved in 20ul solvent A and 15 µL were injected. Samples were trapped on a C18 guard-desalting column (Acclaim PepMap 100, 75µm x 2 cm, nanoViper, C18, 5 µm, 100Å), and separated on a 50 cm long C18 column (Easy spray PepMap RSLC, C18, 2 µm, 100Å, 75 µm x 50 cm). The nano capillary solvent A was 95% water, 5%DMSO, 0.1% formic acid; and solvent B was 5% water, 5% DMSO, 95% acetonitrile, 0.1% formic acid. At a constant flow of 0.25 µl $min^{-1}$, the curved gradient went from 6-8% B up to 40% B in each fraction in a dynamic range of gradient length (**Sup. Table 2**), followed by a steep increase to 100% B in 5 min. FTMS master scans with 60,000 resolution (and mass range 300-1500 m/z) were followed by data-dependent MS/MS (30 000 resolution) on the top 5 ions using higher energy collision dissociation (HCD) at 30% normalized collision energy. Precursors were isolated with a 2 m/z window. Automatic gain control (AGC) targets were $1e^6$ for MS1 and $1e^5$ for MS2. Maximum injection times were 100 ms for MS1 and 100 ms for MS2. The entire duty cycle lasted ~2.5 s. Dynamic exclusion was used with 30 s duration. Precursors with unassigned charge state or charge state 1 were excluded. An underfill ratio of 1% was used.

Orbitrap raw MS/MS files were converted to mzML format using msConvert from the ProteoWizard tool suite.(*25*) Spectra were then searched using MSGF+ (v. 10072)(*26*) and Percolator (v2.08)(*27*), where search results from 8 subsequent fraction were grouped for Percolator target/decoy analysis. All searches were done against the human protein subset of Ensembl 75 in the Galaxy platform(*28*). MSGF+ settings included precursor mass tolerance of 10 ppm, fully-tryptic peptides, maximum peptide length of 50 amino acids and a maximum charge of 6. Fixed modifications were TMT-10plex on lysines and peptide N-termini, and carbamidomethylation on cysteine residues, a variable modification was used for oxidation on methionine residues.

Quantification of TMT-10plex reporter ions was done using OpenMS project's IsobaricAnalyzer (v2.0)(*29*). PSMs found at 1% FDR (false discovery rate) were used to infer gene identities. Protein quantification by TMT 10-plex reporter ions was calculated using TMT PSM ratios to the entire sample set (all 10 TMT-channels) and normalized to the sample median. The median PSM TMT reporter ratio from peptides unique to a gene symbol was used for quantification. Protein false discovery rates were calculated using the picked-FDR method using gene symbols as protein groups and limited to 1% FDR.(*30*) Fold changes were recalculated between individual MDS samples and the average expression in NBM samples to facilitate the identification of differentially expressed proteins (DEP). DEP were matched against DEG from the whole-transcript RNAseq dataset and analysed through gene list enrichment analysis as described above.

**Fixed-cell immunofluorescence**
MitoTracker® Deep Red FM (Thermo Fisher Scientific) was used as a mitochondrial selective probe to stain live MACS-separated cells for 30 min at 4ºC. All washes were carried out in PBSA (PBS+0.2% bovine serum albumin [BSA, Merck]). After the initial MitoTracker staining, cells were washed twice, cytospun onto glass slides and fixed in 4% paraformaldehyde for 10 min at room temperature (RT). Once fixed, cells were washed twice and permeabilized in 0.1% Triton X-100 (Merck) for 5 min at RT. Blocking was performed through incubation in PBS + 2% BSA for 1 h at RT. Cells were washed twice and incubated with primary anti-Ki67 for 1 h at RT. The cells were then washed twice and incubated for 1 h at RT with Alexa Fluor-488 Goat anti-Rabbit IgG, washed twice and finally counterstained with PBS + 0.1 µg/mL DAPI for 10 min at RT. Coverslips were mounted and left to dry at 4ºC for 24 h. Confocal images were acquired using an Olympus IX71 inverted microscope (Olympus Life Science) with 63x/1.4 NA oil-immersion lens and NIS-Elements Basic Research v. 3.2 (Nikon, Japan) image acquisition software, and processed using Fiji v. 2.3.0/1.53f51.(*31*)

### Supplemental Tables

#### Supplemental table 1 – Bulk RNAseq quality control metrics

| | Total dataset metrics | | | | | | Average per cell type | | | | |
|---|---|---|---|---|---|---|---|---|---|---|---|
| | Min | Q1 | Median | Mean | Q3 | Max | CD34+ HSPCs | GPA erythroblasts | CD71+ Retics | Ring sideroblasts | Siderocytes |
| Total sequenced reads (Million) | 21.6 | 32.3 | 37.4 | 46.2 | 56.8 | 127.6 | 56.9 | 36 | 29.8 | 40.8 | 55.1 |
| RNA Integrity number (RIN) | 7.0 | 7.8 | 8.4 | 8.5 | 9.1 | 10.0 | 8.7 | 8.2 | 7.9 | 9.1 | 8.5 |
| Quality-trimmed reads (%) | 0.2 | 0.2 | 0.2 | 0.3 | 0.3 | 0.9 | 0.3 | 0.2 | 0.3 | 0.4 | 0.2 |
| Input read length (bp) | 200 | 262 | 266 | 262 | 268 | 278 | 255 | 267 | 268 | 263 | 262 |
| Uniquely mapped reads (Million) | 4.7 | 10.9 | 18 | 20.3 | 26.5 | 67.7 | 31 | 16.3 | 7.0 | 15.1 | 13.2 |
| Uniquely mapped reads (%) | 21.8 | 29.3 | 46 | 42.4 | 51.6 | 65.6 | 54.2 | 44.8 | 23.3 | 35.2 | 24.4 |
| Mapped read length (bp) | 223 | 265 | 268 | 266 | 270 | 278 | 261 | 270 | 272 | 268 | 264 |
| Multimapped reads (Million) | 12 | 17.5 | 20.4 | 23.7 | 23.6 | 69.6 | 23.3 | 18.8 | 21.9 | 23 | 38.8 |
| Multimapped reads (%) | 29.8 | 44.8 | 49.9 | 53.7 | 65.2 | 75.5 | 41.4 | 52.5 | 73.7 | 59.8 | 70.4 |
| Overly multimapped reads (Million) | 0.1 | 0.1 | 0.1 | 0.2 | 0.2 | 1.5 | 0.3 | 0.1 | 0.1 | 0.2 | 0.2 |
| Overly multimapped reads (%) | 0.3 | 0.3 | 0.3 | 0.4 | 0.4 | 2.2 | 0.5 | 0.3 | 0.4 | 0.4 | 0.4 |
| Unmapped reads (Million) | 0.5 | 0.8 | 0.9 | 1.8 | 2.1 | 7.9 | 2.3 | 0.8 | 0.8 | 2.6 | 2.8 |
| Unmapped reads (%) | 1.9 | 2.3 | 2.5 | 3.5 | 2.9 | 11 | 3.8 | 2.3 | 2.6 | 4.6 | 4.7 |

#### Supplemental table 2 – RNA/protein GO enrichment results comparing *SF3B1*$^{mt}$ RS vs. healthy donor EB.

| Group | RNA ↑ Protein ↑ | | RNA ↓ Protein ↑ | | RNA ↓ Protein ↓ | | RNA ↑ Protein ↓ | |
|---|---|---|---|---|---|---|---|---|
| | GO BP Term | Adj. P-value | GO BP Term | Adj. P-value | GO BP Term | Adj. P-value | GO BP Term | Adj. P-value |
| Non-redundant sig. enriched GO BP terms | Ubiquitin proteolysis | $1.26 \times 10^{-7}$ | Metabolic process | $7.68 \times 10^{-14}$ | Cell cycle | $9.19 \times 10^{-35}$ | Chromatin organization | $1.99 \times 10^{-5}$ |
| | Autophagy | $2.00 \times 10^{-6}$ | Amino acid metabolism | $1.39 \times 10^{-5}$ | Leukocyte activation | $7.63 \times 10^{-33}$ | Histone modification | $1.99 \times 10^{-5}$ |
| | Response to hypoxia | $6.91 \times 10^{-6}$ | Purine biosynthesis | $2.47 \times 10^{-5}$ | DNA replication | $1.14 \times 10^{-19}$ | Regulation of transcription | $1.50 \times 10^{-4}$ |
| | Translation | $1.20 \times 10^{-4}$ | Glycosyl compound metabolism | $3.19 \times 10^{-5}$ | Vesicle-mediated transport | $9.46 \times 10^{-19}$ | Organelle organization | $8.60 \times 10^{-3}$ |
| | Response to ER stress | $5.20 \times 10^{-4}$ | Mitochondrial targeting | $1.80 \times 10^{-3}$ | Chromosome organization | $4.46 \times 10^{-18}$ | Regulation of gene expression | $1.80 \times 10^{-3}$ |
| | Response to ROS | $3.90 \times 10^{-3}$ | Sulfur compound biosynthesis | $1.00 \times 10^{-2}$ | Cytoskeleton organization | $1.08 \times 10^{-12}$ | Protein metabolic process | $1.60 \times 10^{-2}$ |
| | Glucose homeostasis | $5.60 \times 10^{-3}$ | | | Hemopoiesis | $1.95 \times 10^{-12}$ | | |
| | Mitophagy | $6.30 \times 10^{-3}$ | | | Cell death | $6.80 \times 10^{-4}$ | | |
| | Iron ion transport | $1.07 \times 10^{-2}$ | | | Cell-cell adhesion | $1.60 \times 10^{-3}$ | | |

### Supplemental table 3 – Antibodies and fluorescent markers

| Antibody/Reagent | ID | Conjugate | Species | Clone | Source |
|---|---|---|---|---|---|
| **Rabbit IgG (H+L)** | A-11008 | Alexa Fluor-488 | Goat | Polyclonal | Thermo Fisher Scientific |
| **CD44** | 338813 | Alexa Fluor-700 | Mouse | BJ18 | Biolegend |
| **CD34** | 555821 | APC | Mouse | 581 | BD Biosciences |
| **CD49d (ITGA4)** | 130-123-797 | APC | Mouse | MZ18-24A9 | Miltenyi Biotec |
| **CD71 (TFRC)** | 334109 | APC-Cy7 | Mouse | CY1G4 | Biolegend |
| **CD235a (GPA)** | 13-9987-82 | Biotin | Mouse | HIR2 | eBiosciences |
| **CD45RA** | 404-0458-42 | BV421 | Mouse | HI100 | Invitrogen |
| **Ki-67 (MKI67)** | 350506 | BV421 | Mouse | Ki-67 | Biolegend |
| **CD233 (Band 3)** | 08-9439-3 | FITC | Mouse | BRIC 6 | Emelca Bioscience |
| **CD71 (TFRC)** | 561939 | FITC | Mouse | M-A712 | BD Biosciences |
| **CD235a (GPA)** | 349106 | PE | Mouse | HI264 | Biolegend |
| **CD90** | 328110 | PE | Mouse | 5E10 | Biolegend |
| **CD3** | 300410 | PE-Cy5 | Mouse | UCHT1 | Biolegend |
| **CD4** | 300510 | PE-Cy5 | Mouse | RPA-T4 | Biolegend |
| **CD7** | 343110 | PE-Cy5 | Mouse | CD7-6B7 | Biolegend |
| **CD8** | 301010 | PE-Cy5 | Mouse | RPA-T8 | Biolegend |
| **CD10** | 312206 | PE-Cy5 | Mouse | HI10a | Biolegend |
| **CD11b** | 301308 | PE-Cy5 | Mouse | ICRF44 | Biolegend |
| **CD14** | A07765 | PE-Cy5 | Mouse | RMO52 | Beckman Coulter |
| **CD19** | 302210 | PE-Cy5 | Mouse | HIB19 | Biolegend |
| **CD20** | 302308 | PE-Cy5 | Mouse | 2H7 | Biolegend |
| **CD56** | 555517 | PE-Cy5 | Mouse | B159 | BD Biosciences |
| **CD123** | 306010 | PE-Cy7 | Mouse | 6H6 | Biolegend |
| **CD38** | 303538 | PE-Dazzle594 | Mouse | HIT2 | Biolegend |
| **CD235a (GPA)** | 306627 | TotalSeq-B | Mouse | HIR2 | Biolegend |
| **Ki67** | 12202 | Unconjugated | Rabbit | D3B5 | Cell Signalling |
| **Streptavidin** | 561419 | V500 | ----------- | ----------- | BD Biosciences |
| **7AAD** | 00-6993-50 | ----------- | ----------- | ----------- | eBiosciences |
| **Hoechst 33342** | B2261 | ----------- | ----------- | ----------- | Sigma-Aldrich |
| **MitoTracker Deep Red FM** | M22426 | ----------- | ----------- | ----------- | Thermo Fisher Scientific |
| **Thiazole Orange** | 390062 | ----------- | ----------- | ----------- | Sigma-Aldrich |
| **DAPI** | D9542 | ----------- | ----------- | ----------- | Sigma-Aldrich |
| **DRAQ5** | 62251 | ----------- | ----------- | ----------- | Thermo Fisher |

#### Supplemental table 4 – Gradient length of HiRIEF fractions

| Sample | Gradient length | Sample | Gradient length |
|---|---|---|---|
| fraction_01 | 50 min | fraction_37 | 50 min |
| fraction_02 | 50 min | fraction_38 | 70 min |
| fraction_03 | 70 min | fraction_39 | 70 min |
| fraction_04 | 70 min | fraction_40 | 50 min |
| fraction_05 | 90 min | fraction_41 | 50 min |
| fraction_06 | 90 min | fraction_42 | 50 min |
| fraction_07 | 110 min | fraction_43 | accumulate in trap column |
| fraction_08 | 110 min | fraction_44 | accumulate in trap column |
| fraction_09 | 110 min | fraction_45 | accumulate in trap column |
| fraction_10 | 110 min | fraction_46 | accumulate in trap column |
| fraction_11 | 110 min | fraction_47 | accumulate in trap column |
| fraction_12 | 90 min | fraction_48 | accumulate in trap column |
| fraction_13 | 90 min | fraction_49 | accumulate in trap column |
| fraction_14 | 90 min | fraction_50 | 50 min |
| fraction_15 | 90 min | fraction_51 | 70 min |
| fraction_16 | 70 min | fraction_52 | 70 min |
| fraction_17 | 50 min | fraction_53 | accumulate in trap column |
| fraction_18 | 50 min | fraction_54 | accumulate in trap column |
| fraction_19 | 50 min | fraction_55 | accumulate in trap column |
| fraction_20 | accumulate in trap column | fraction_56 | accumulate in trap column |
| fraction_21 | accumulate in trap column | fraction_57 | accumulate in trap column |
| fraction_22 | accumulate in trap column | fraction_58 | accumulate in trap column |
| fraction_23 | accumulate in trap column | fraction_59 | accumulate in trap column |
| fraction_24 | accumulate in trap column | fraction_60 | accumulate in trap column |
| fraction_25 | accumulate in trap column | fraction_61 | accumulate in trap column |
| fraction_26 | accumulate in trap column | fraction_62 | accumulate in trap column |
| fraction_27 | 50 min | fraction_63 | accumulate in trap column |
| fraction_28 | 50 min | fraction_64 | 50 min |
| fraction_29 | 70 min | fraction_65 | 50 min |
| fraction_30 | 50 min | fraction_66 | 70 min |
| fraction_31 | 50 min | fraction_67 | accumulate in trap column |
| fraction_32 | accumulate in trap column | fraction_68 | accumulate in trap column |
| fraction_33 | accumulate in trap column | fraction_69 | accumulate in trap column |
| fraction_34 | accumulate in trap column | fraction_70 | accumulate in trap column |
| fraction_35 | accumulate in trap column | fraction_71 | 50 min |
| fraction_36 | 50 min | fraction_72 | 50 min |

#### Supplemental figures

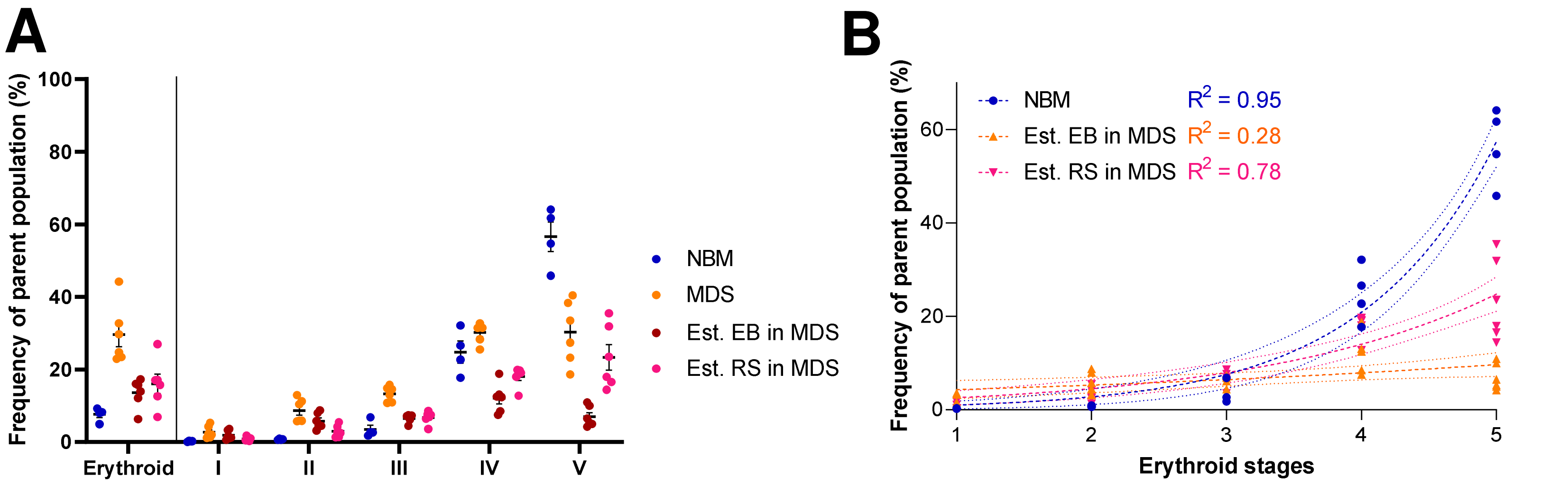

##### Supplemental figure 1

**A)** Mean (±SEM) cell population frequencies within the respective parent populations in FC, quantified in normal bone marrow (NBM) donors ($n^{NBM}$ = 4) and MDS-RS patients ($n^{MDS}$ = 6). Erythroid cells are quantified within the singlet population, EB subsets are quantified within the GPA+ population. Estimated EB and RS contribution to each stage is calculated through the observed RS frequencies per stage (**Figure 1C**).

**B)** Progression through erythroid stages for normal EB and estimated MDS subpopulations is fitted to theoretical exponential growth curves.

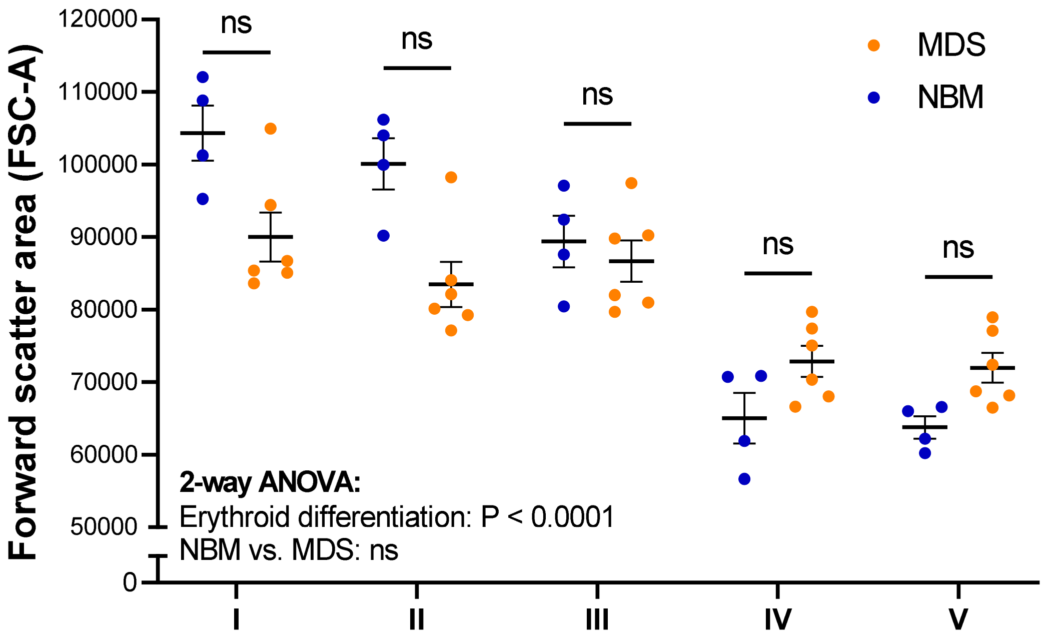

**Supplemental figure 2**

Mean (± SEM) forward scatter area (FSC-A, cell size surrogate) per EB subset ($n^{NBM}$ = 4, $n^{MDS\text{-}RS}$ = 6).

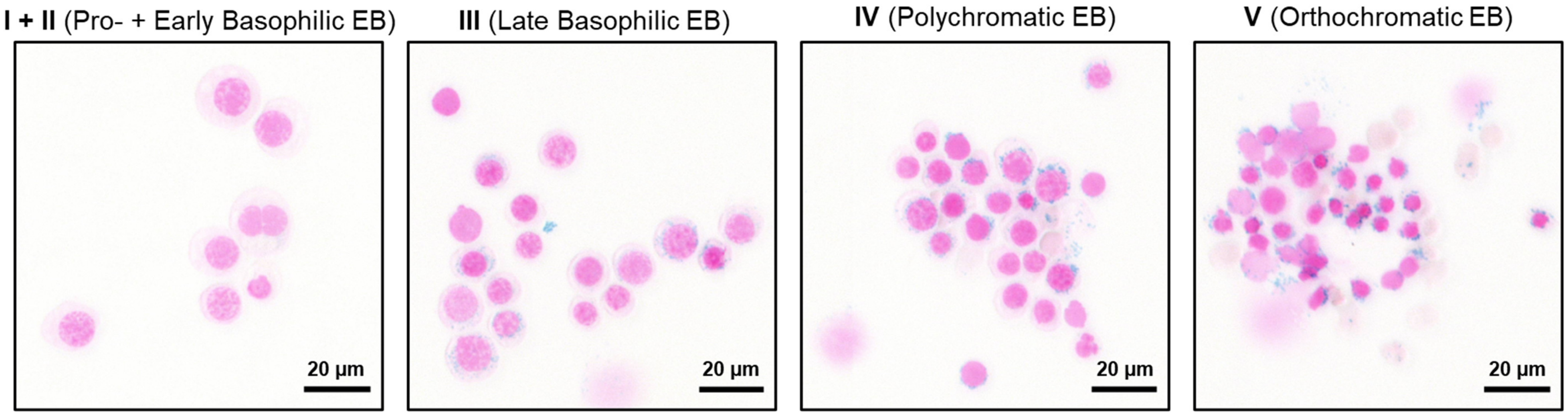

**Supplemental figure 3**

Representative micrographs of FACS-isolated MDS-RS EB subset cytospins stained with Prussian Blue. Iron granules stain blue, hemoglobin brown and DNA pink. Scale bars: 20 µm.

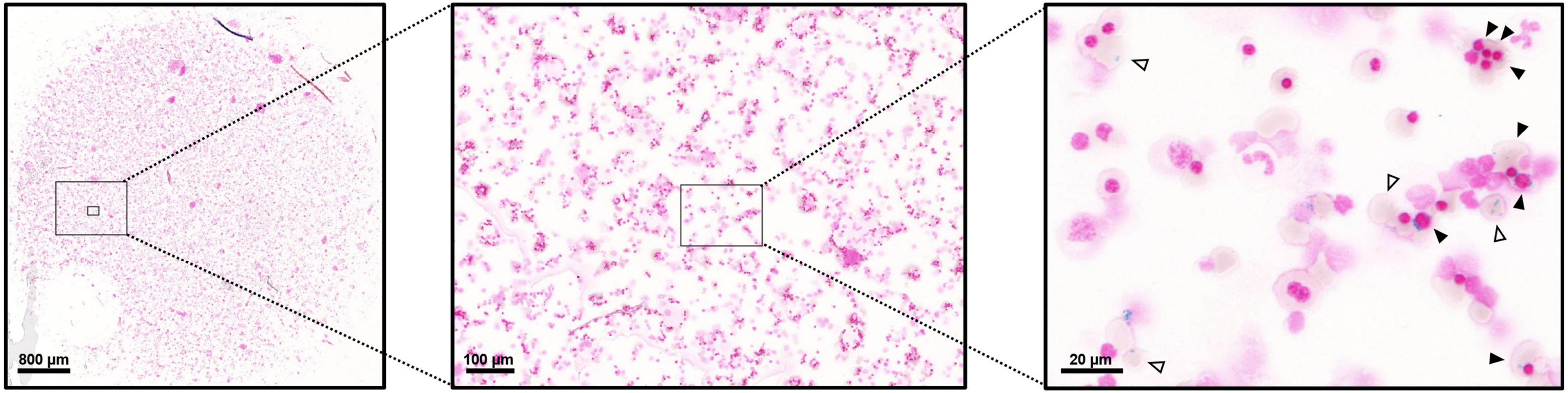

**Supplemental figure 4**

Representative micrograph of a cytospin obtained from 28-day culture of MDS-RS MNC, stained with Prussian Blue. Filled arrows indicate ring sideroblasts, outlined arrows indicate siderocytes. Iron granules stain blue, hemoglobin brown and DNA pink. Scale bars: 800 µm, 100 µm and 20 µm.

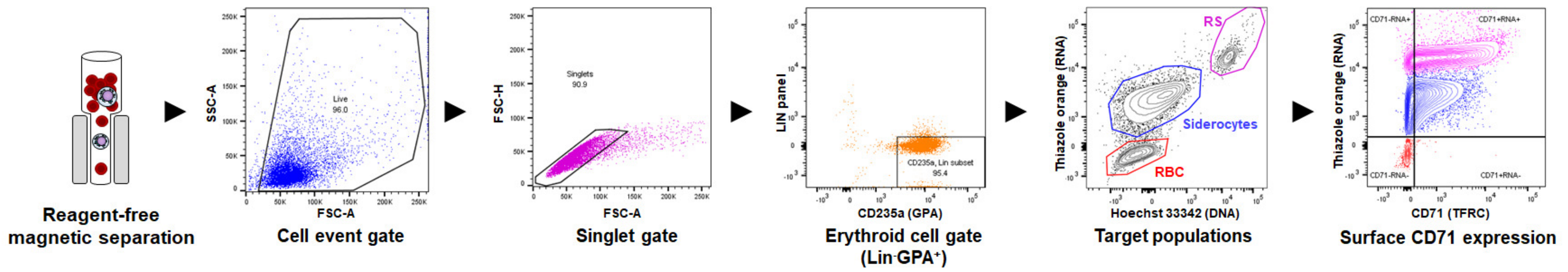

#### Supplemental figure 5

Complete sorting strategy comprising gating steps to isolate purified ring sideroblasts and siderocytes from the high density cell layer after initial MACS enrichment, including surface CD71 quantification.

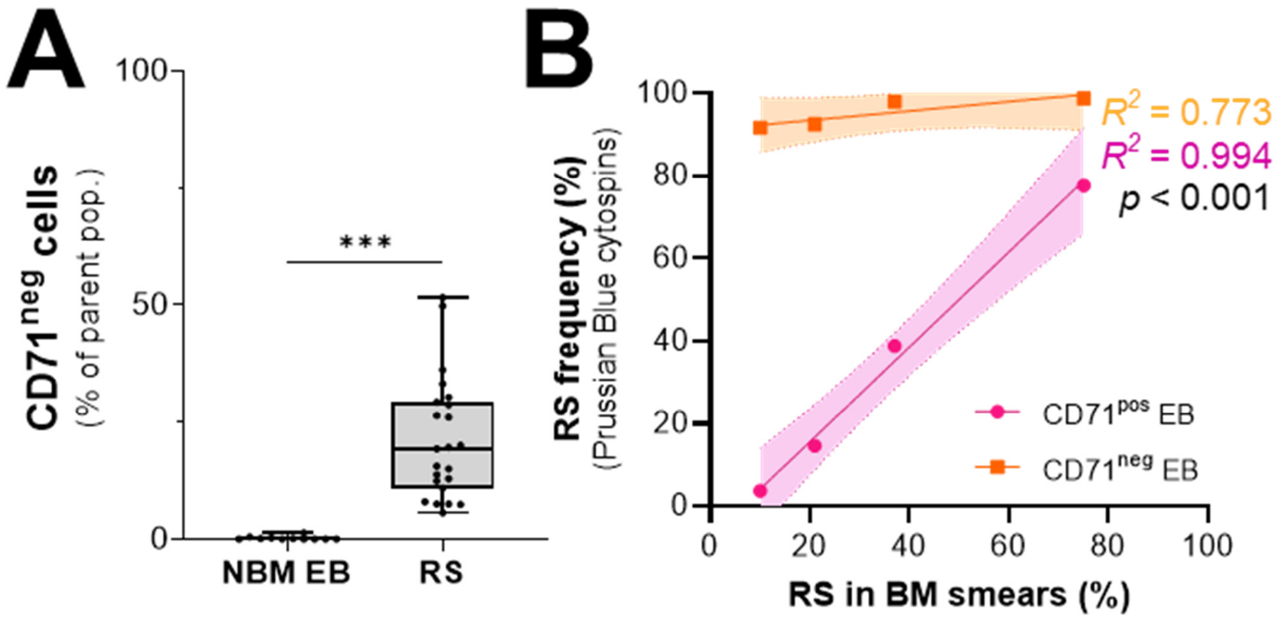

**Supplemental figure 6**

**A)** Percentage of CD71-negative cells in flow cytometry evaluation among erythroblasts from normal bone marrow donors (NBM) (n = 11) and *SF3B1*$^{mt}$ RS (n = 23). **B)** Correlation of clinically-determined RS percentages and RS frequencies in cytospins obtained from FACS purification of CD71-expressing and non-expressing erythroblasts (Lin-GPA+DNA+).

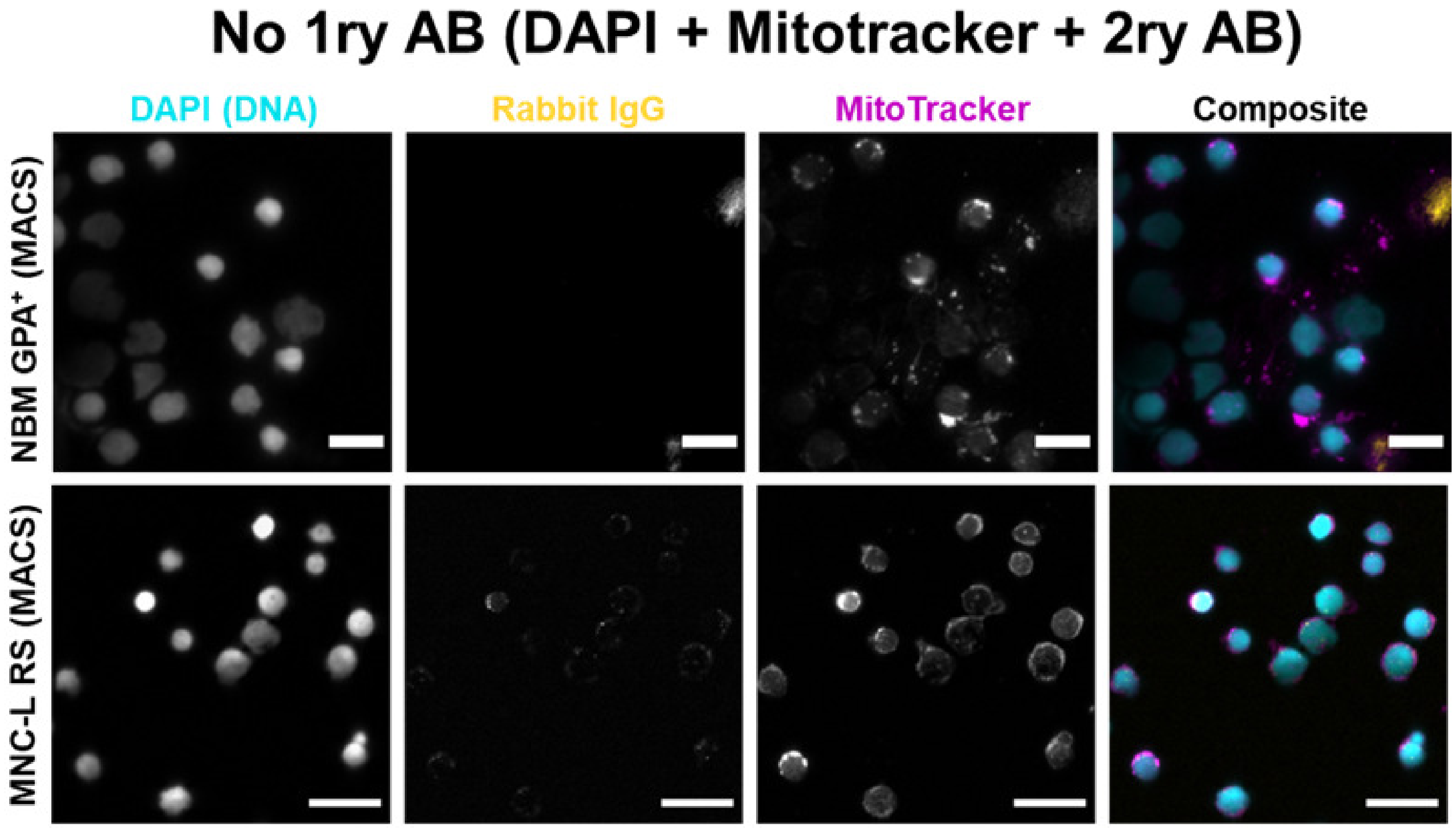

**Supplemental figure 7**

Immunofluorescence displaying Ki-67 signal in NBM GPA$^{+}$ and an RS isolate (MNC-L, average RS isolation purity ≈ 90%) with only secondary antibody added and no primary anti-Ki67. The sample was thus co-labeled for DAPI (DNA marker; cyan), anti-Rabbit IgG (yellow) and MitoTracker Deep Red (mitochondrial marker; magenta) and imaged via widefield microscopy. Individual channels are presented in greyscale, with a composite image displaying all three markers in the indicated colors. Scale bars = 20 µm.

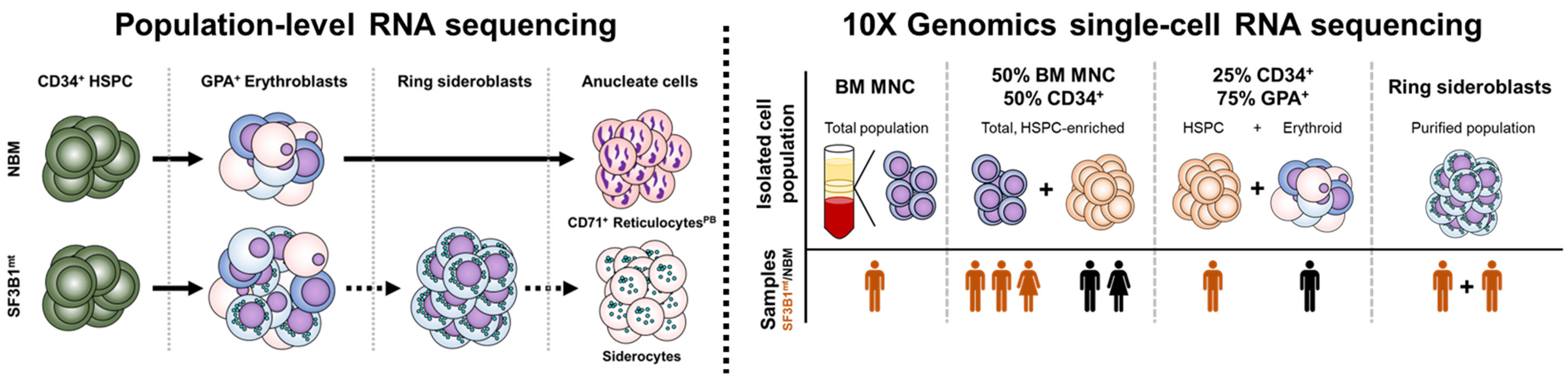

**Supplemental figure 8**

Experimental design of whole-transcript population-level RNA sequencing (left) and 10X Genomics single-cell RNA sequencing (right) encompassing NBM donors and SF3B1mt MDS-RS patients.

Population-level RNAseq: Hematopoietic stem cell progenitor (HSPC) samples ($n^{NBM}$ = 7, $n^{MDS}$ = 6), mixed erythroblast samples ($n^{NBM}$ = 4, $n^{MDS}$ = 5) and peripheral blood reticulocyte samples ($Ret^{PB}$) ($n^{NBM}$ = 4) were obtained through MACS with CD34, GPA and CD71-specific antibodies, respectively. RS ($n^{MDS}$ = 4) and siderocytes ($n^{MDS}$ = 4) were obtained through M+FACS.

scRNAseq: 3 NBM and 5 MDS-RS patients were assayed over two separate 10X experiments and integrated during analysis. In total, one BM MNC sample ($n^{MDS}$ = 1), five HSPC-enriched samples (1:1 BM MNCs and $CD34^{+}$-enriched cells, $n^{NBM}$ = 2, $n^{MDS}$ = 3), two HSPC and erythroid-specific samples (1:3 $CD34^{+}$ and $GPA^{+}$-enriched cells, $n^{NBM}$ = 1, $n^{MDS}$ = 1) and one combined RS sample from two patients (purified and mixed 1:1) were processed.

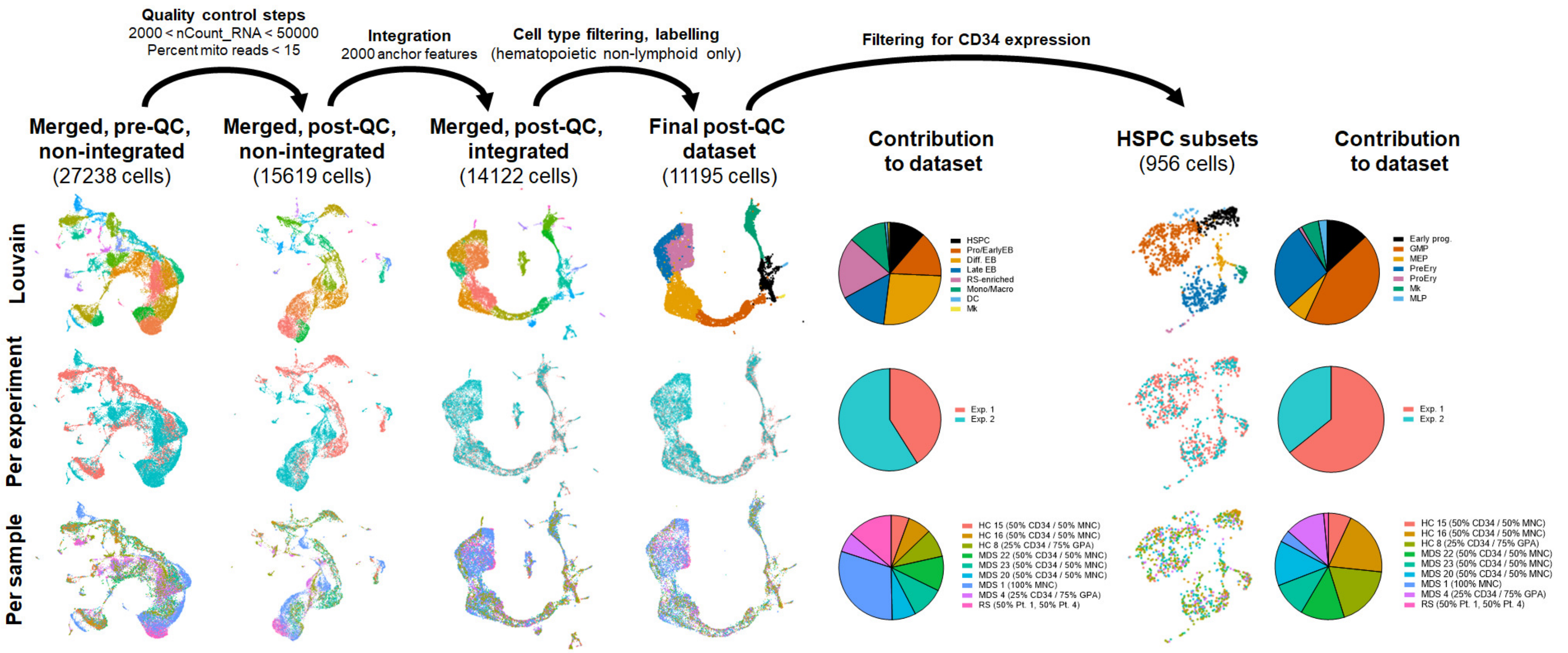

#### Supplemental figure 9

Preprocessing steps for analysis of the scRNAseq dataset, including cell type, 10X experiment and donor/patient sample contribution to the dataset at both the total level and specifically in cells expressing CD34 transcripts.

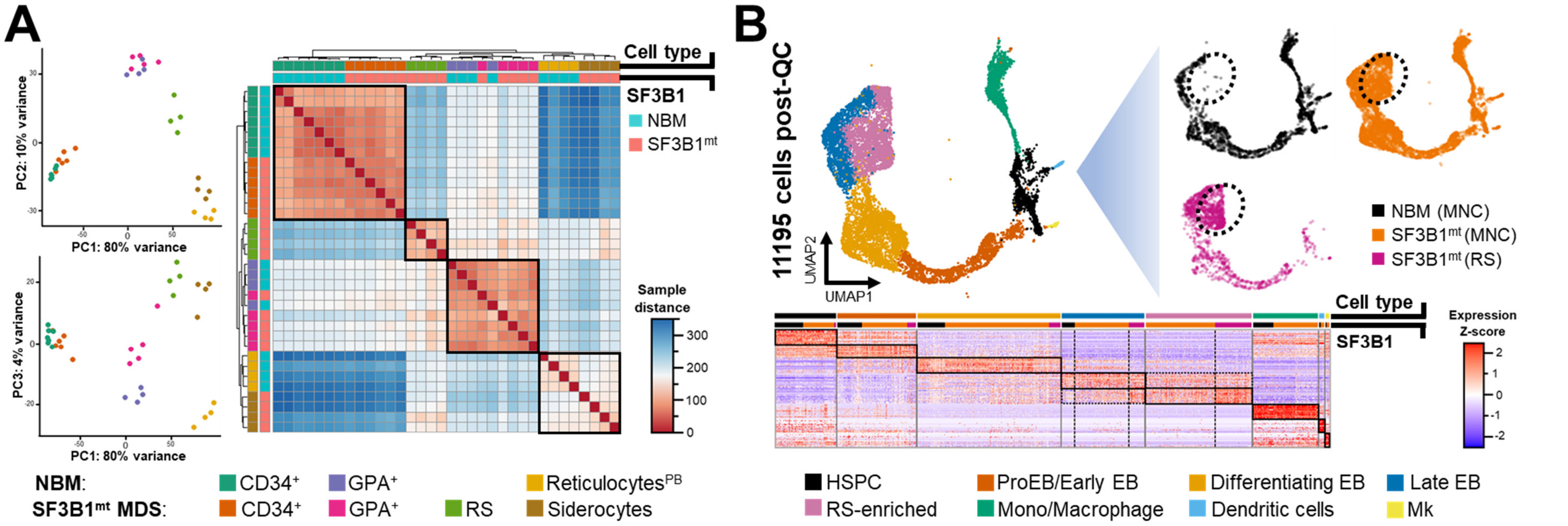

**Supplemental figure 10**

Global overviews of **A)** whole-transcript population-level RNAseq, including an inter-sample distance heatmap to visualize cell type and subset separation and **B)** scRNAseq, including a gene expression heatmap of differentially expressed genes per major transcriptomically identifiable cell type and according to SF3B1 background (NBM, MDS-RS MNC from *SF3B1*$^{mt}$ patients and *SF3B1*$^{mt}$ RS).

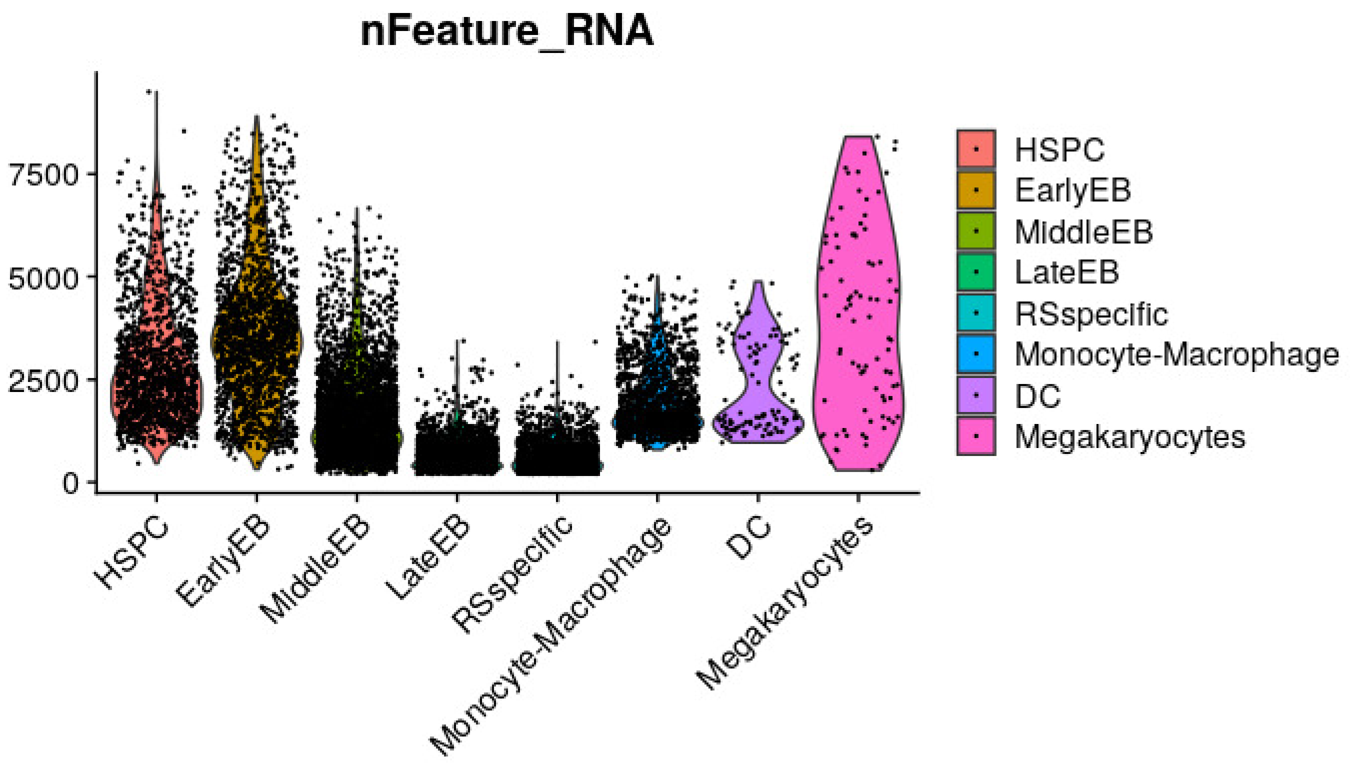

**Supplemental figure 11**

Violin plots of the number of genes detected in 10X scRNAseq per cell type after integration and QC steps.

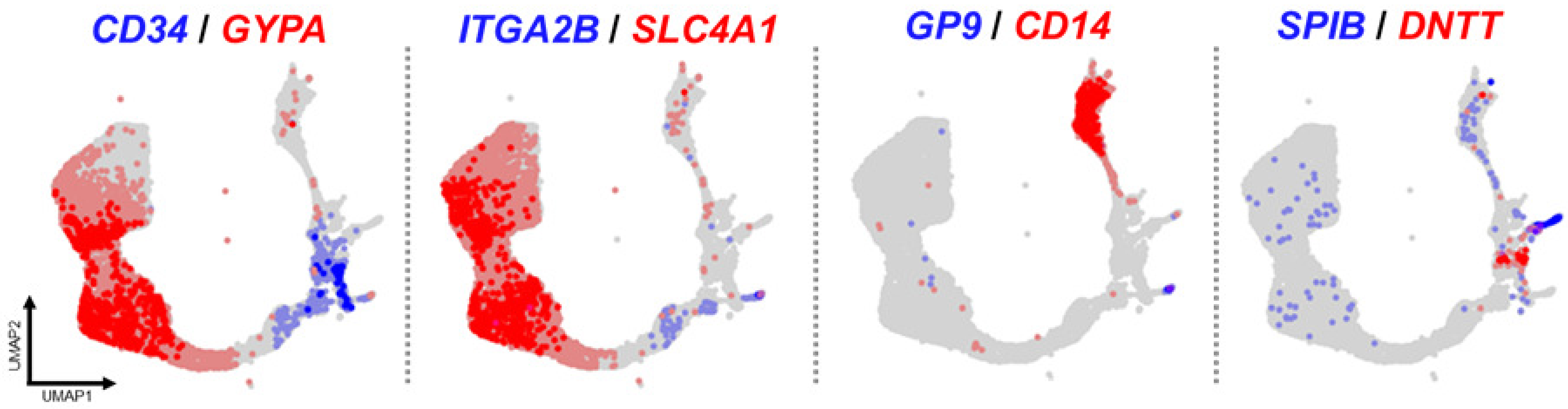

**Supplemental figure 12**

UMAP overlay of the total non-lymphoid hematopoietic dataset, comprising gene expression of selected marker genes (*CD34*: HSPCs and early stage precursors; *GYPA* [Glycophorin A]: differentiating erythroblasts; *ITGA2B* [Integrin α2β]: megakaryocyte-erythroid progenitors, pre-erythroblasts and megakaryocytes; *SLC4A1* [Band 3 / Anion Exchanger 1]: differentiating and late-stage erythroblasts; *GP9* [Glycoprotein IX Platelet]: Megakaryocytes; *CD14*: monocyte-macrophage lineage; *SPIB* [Spi-B Transcription Factor]: dendritic cell lineage; *DNTT* [DNA Nucleotidylexotransferase]: early lymphoid lineage).

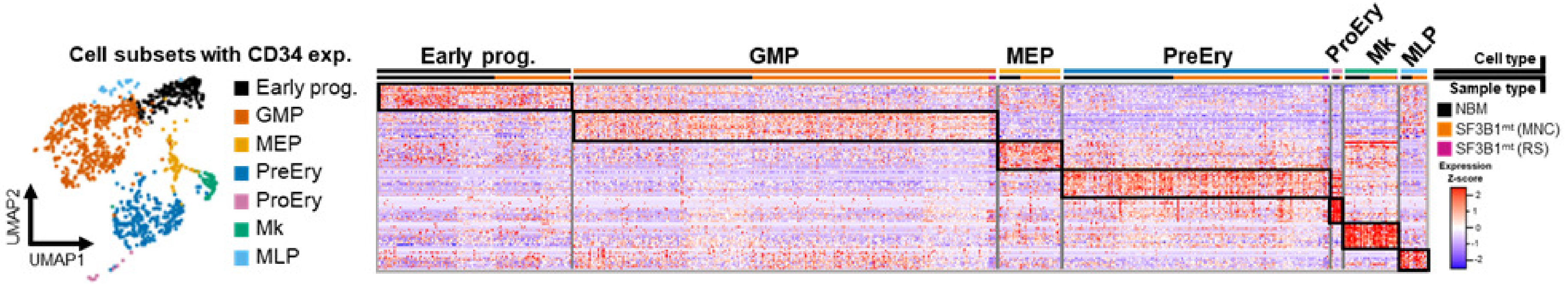

**Supplemental figure 13**

UMAP projection of hematopoietic cells positive for *CD34* RNA expression. Mapping accuracy and cell type annotation was improved through integration of our dataset with the hematopoietic stem cell scRNAseq dataset published by Pellin et al., 2019. The top 50 marker genes for each cluster are displayed in an expression Z-score heatmap (right), where each black box highlights the markers associated with each cluster. The upper bar above the heatmap identifies each cell type as per the UMAP visualization, and cells are further clustered according to sample type, identified by the lower bar. A full list of identified marker genes is provided in **Sup. Data 6**.

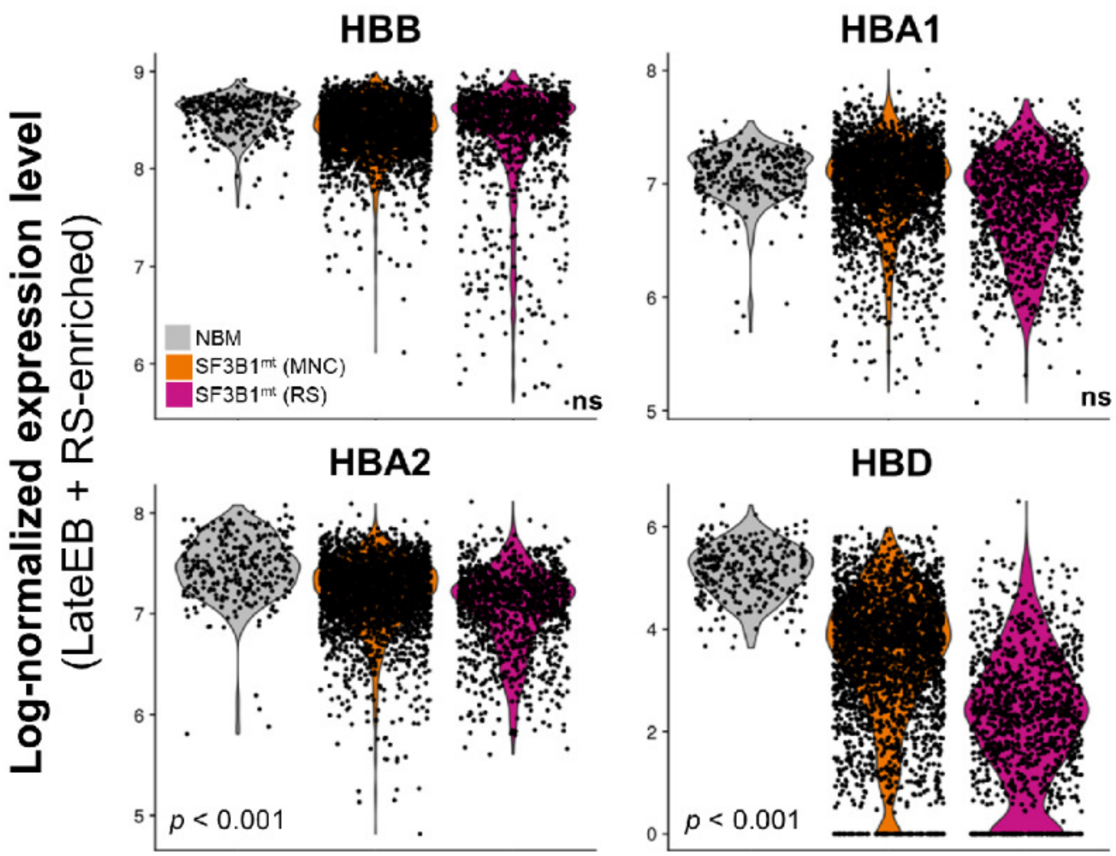

**Supplemental figure 14**

Violin plots of hemoglobin alpha 1, alpha 2, beta and delta chain expression in late erythropoiesis, separated by sample type.

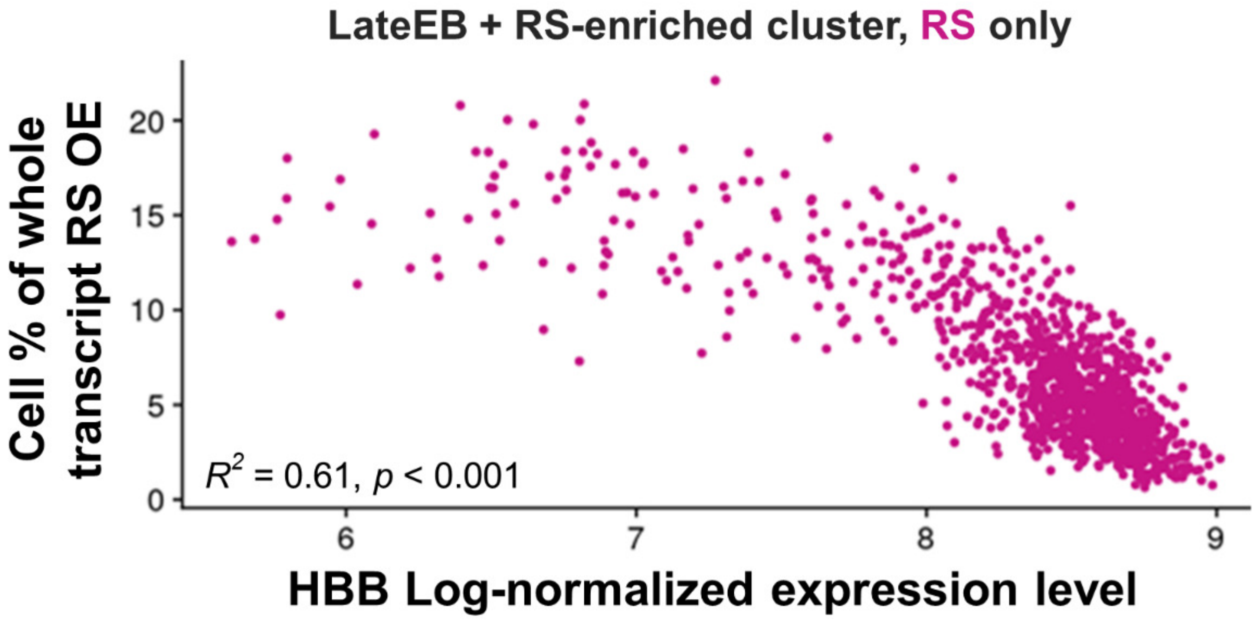

**Supplemental figure 15**

Correlation plot of hemoglobin beta expression and the frequency of RS-signature transcripts in late erythroid RS.

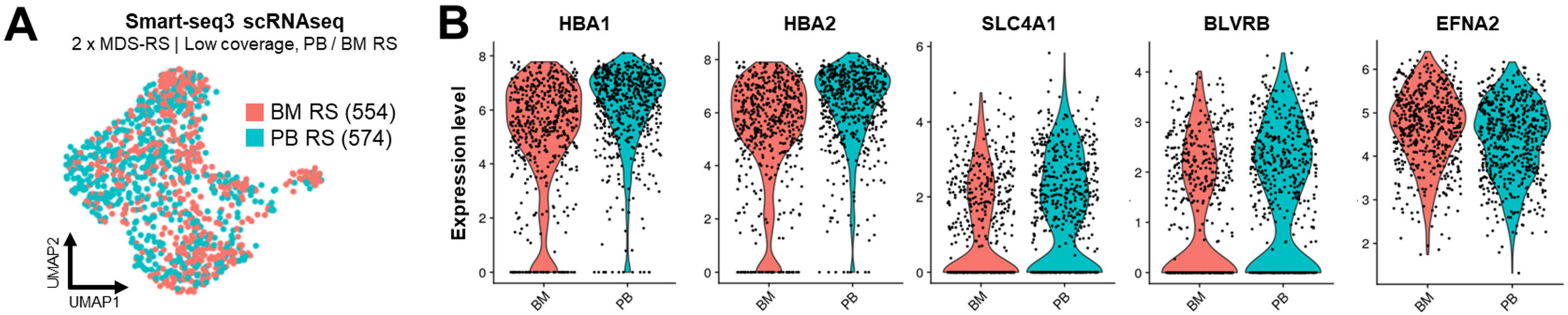

**Supplemental figure 16**

**A)** UMAP visualization of sorted BM and PB RS isolated from 2 patients and analysed through Smart-seq3 (SS3).
**B)** Violin plots displaying selected significant DE erythroid genes comparing BM vs. PB RS.

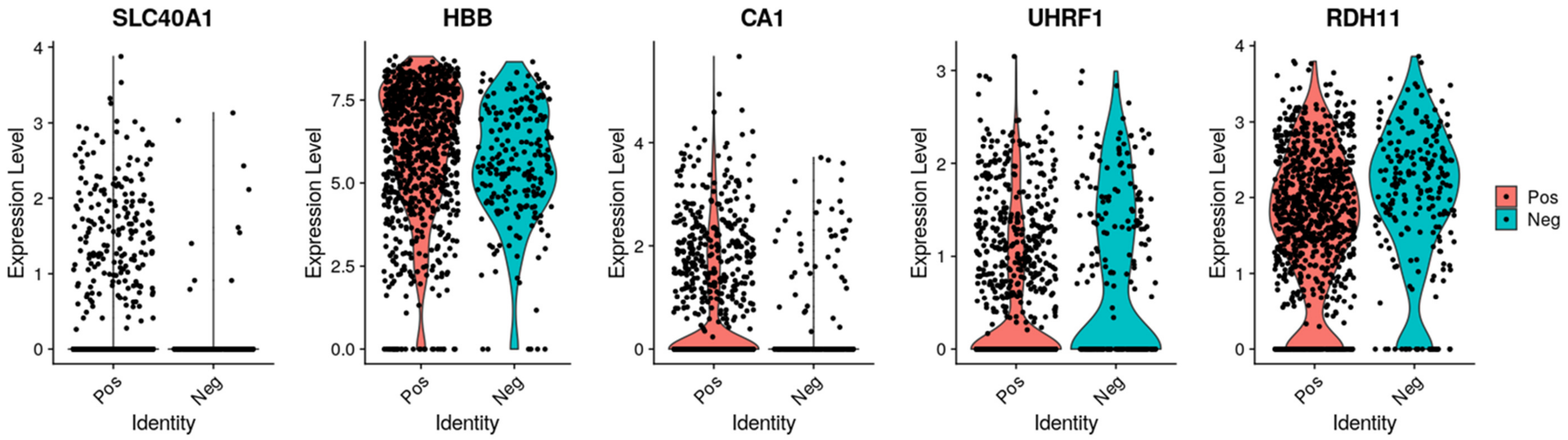

**Supplemental figure 17**

Violin plots comprising several significantly differentially expressed genes comparing CD71-positive RS versus CD71-negative RS (RBC equivalent) in Smart-seq3 RS data.

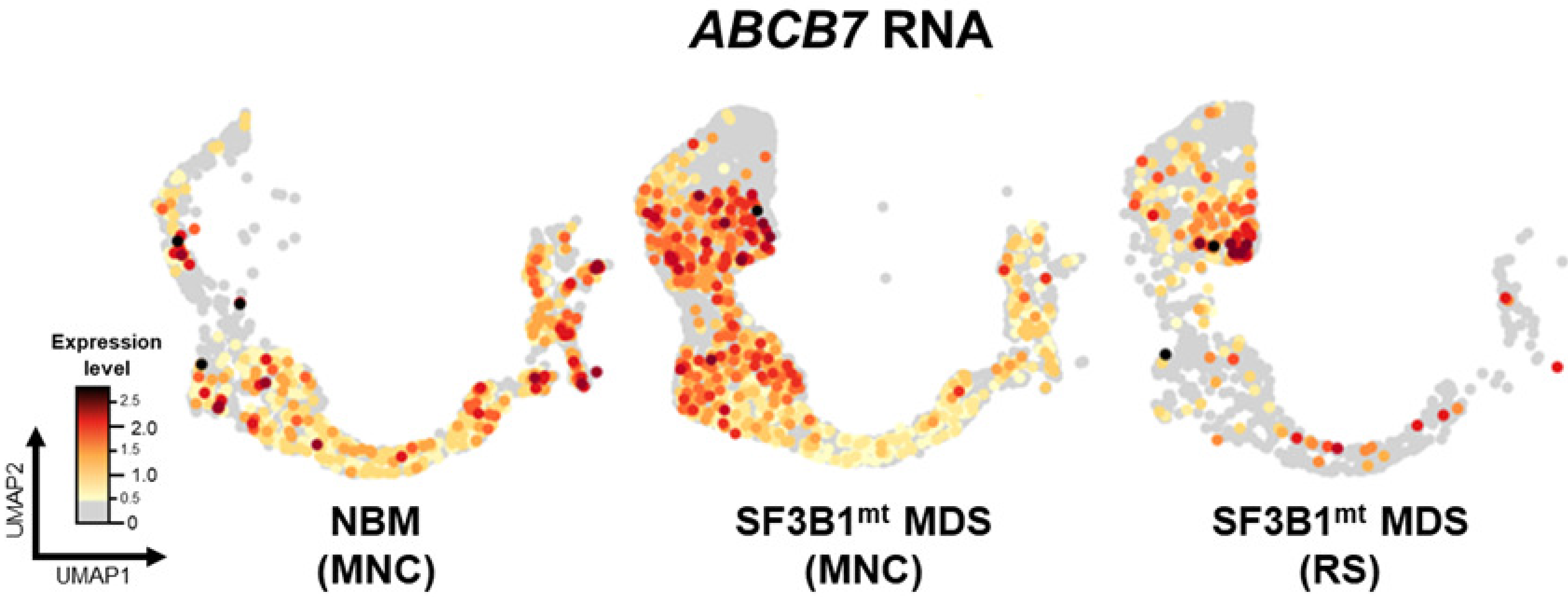

**Supplemental figure 18**

Gene expression of *ABCB7* is overlaid in the HSPC/erythroid UMAP projection, with grey cells displaying no detectable expression and a gradient from light yellow to dark red indicating the level of gene expression per cell. The total dataset is separated by general sample type, specifically NBM (MNC-derived), SF3B1$^{mt}$ MDS (MNC-derived) and SF3B1$^{mt}$ MDS (HD RS).

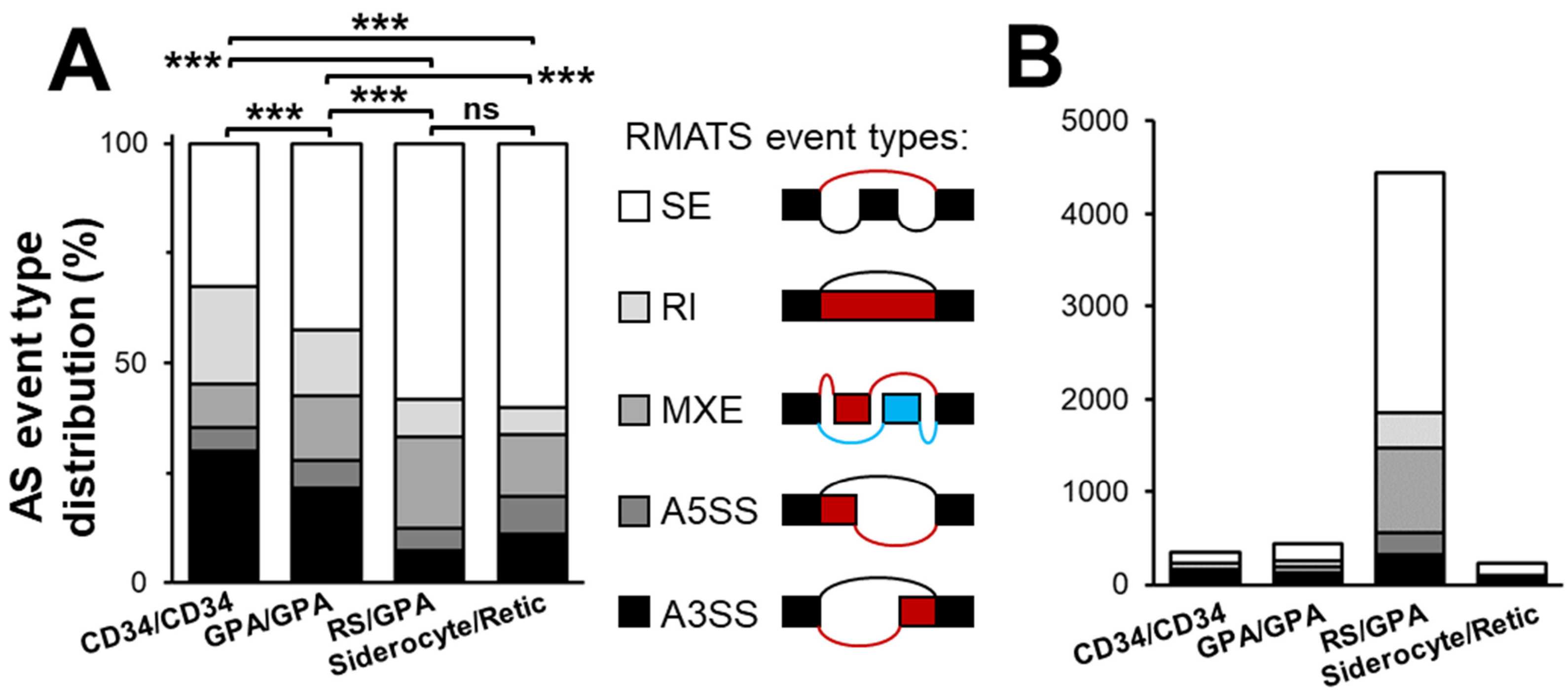

**Supplemental figure 19**

**A)** Frequency and **B)** absolute number of differentially AS events split by rMATS category in each sample group comparison (SE = skipped exon, RI = retained intron, MXE = mutual exon exclusion, A5SS = alternative 5’ splice site, A3SS = alternative 3’ splice site).

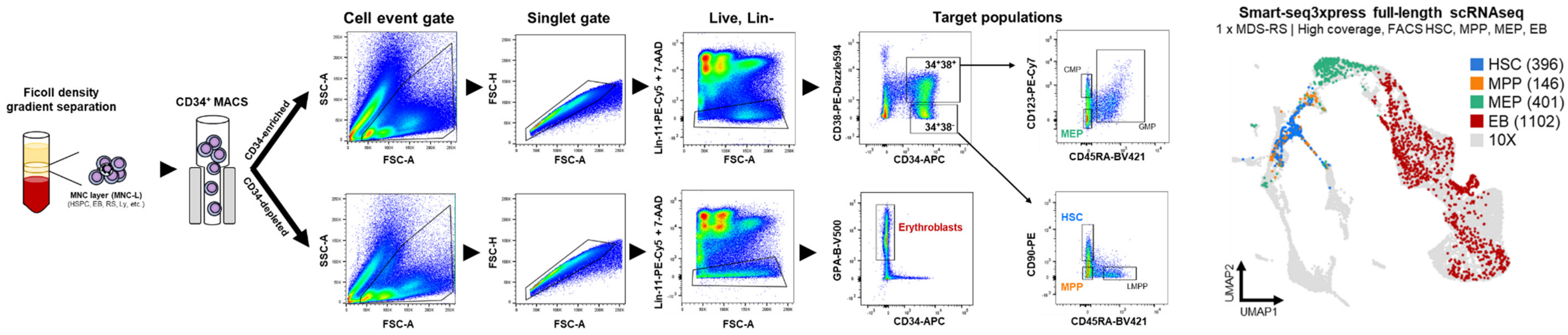

#### Supplemental figure 20

Complete sorting strategy for Smart-seq3xpress, comprising gating steps to isolate HSPC subsets (HSC, MPP, MEP) and erythroblasts after initial CD34$^{+}$ MACS separation. The sorted cell populations are mapped in the integrated 10X/SS3x UMAP plot.

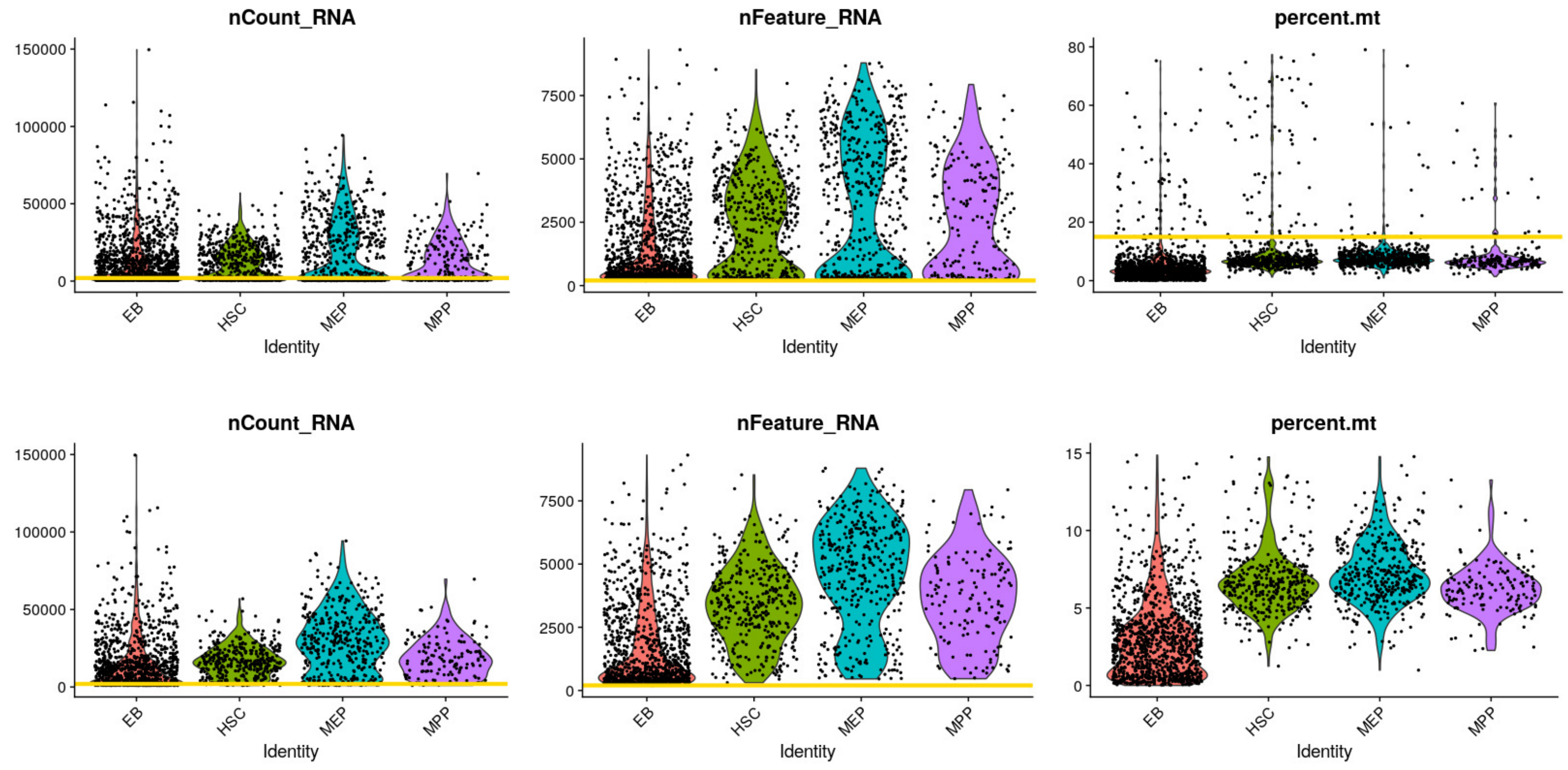

**Supplemental figure 21**

Violin plots of RNA transcript counts per cell type (left), number of genes detected per cell type (center) and percentage of mitochondrial genes per cell type (right), before (top row) and after (bottom row) QC steps. QC cut-offs are indicated with a yellow line.

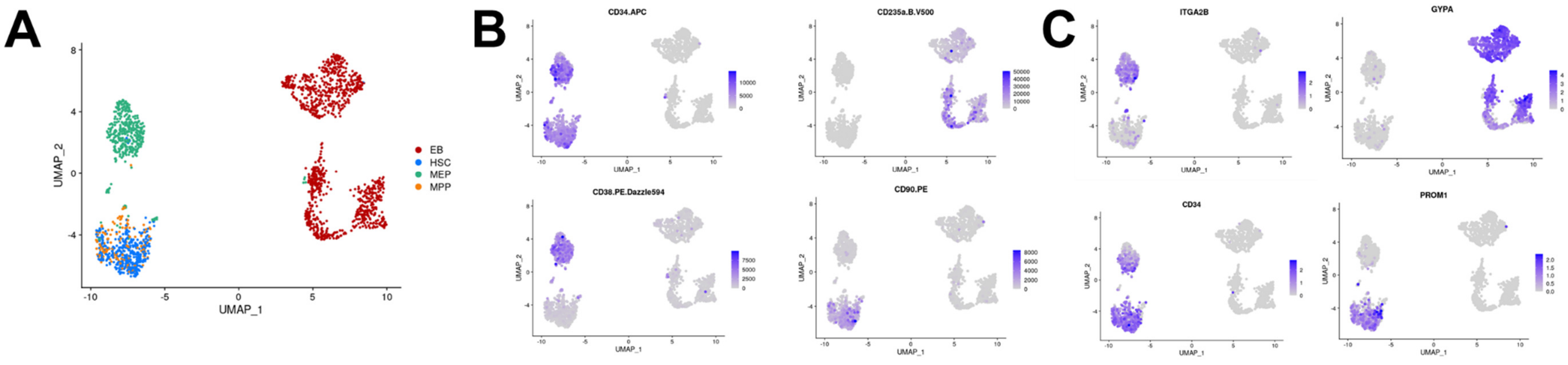

#### Supplemental figure 22

**A)** UMAP overlay of sorted cell subsets analysed with Smart-seq3xpress. B) UMAP overlay of index sorting flow cytometry data on the UMAP plot, with identifying markers CD34, CD38, CD90, CD235a (GPA). C) UMAP overlay of identifying marker gene expression.

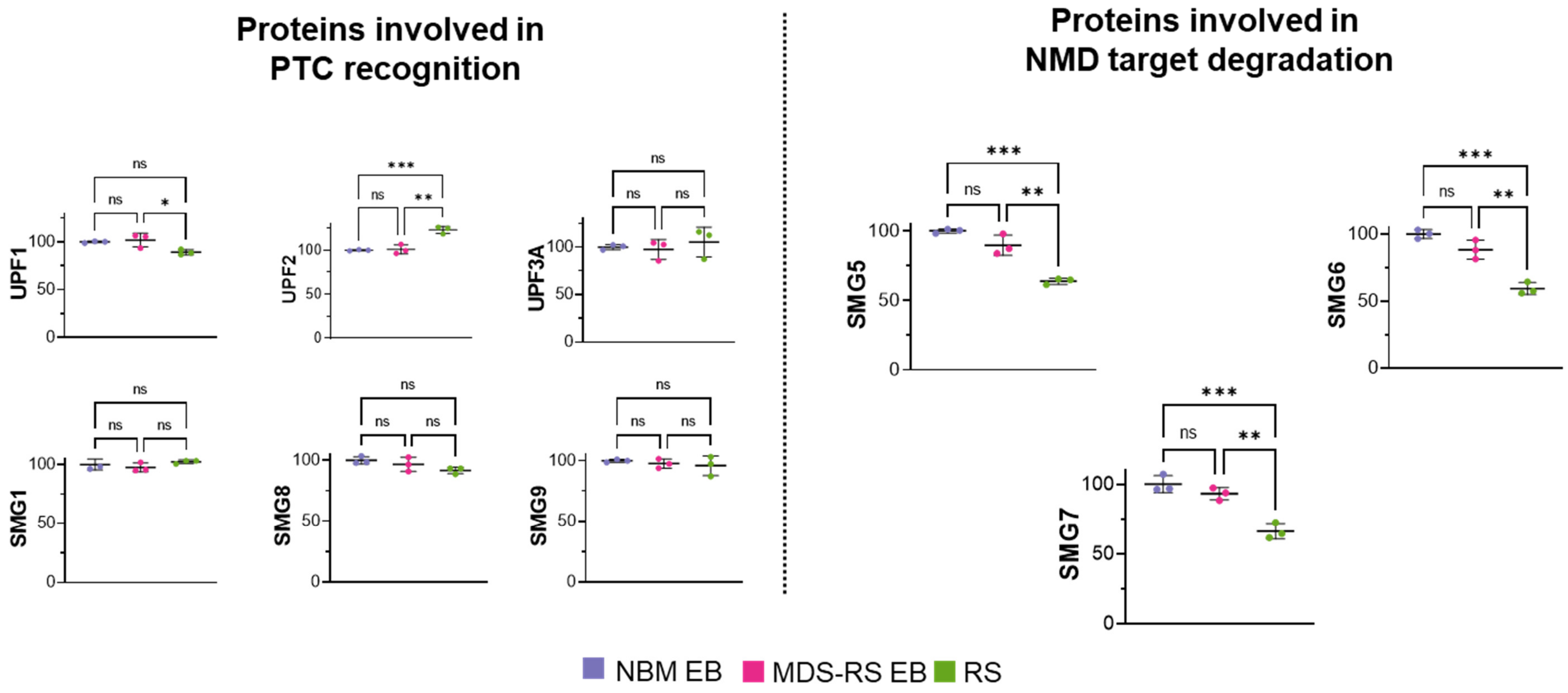

**Supplemental figure 23**

Quantification of NMD pathway-involved proteins in each cell type, separated by major function in NMD.

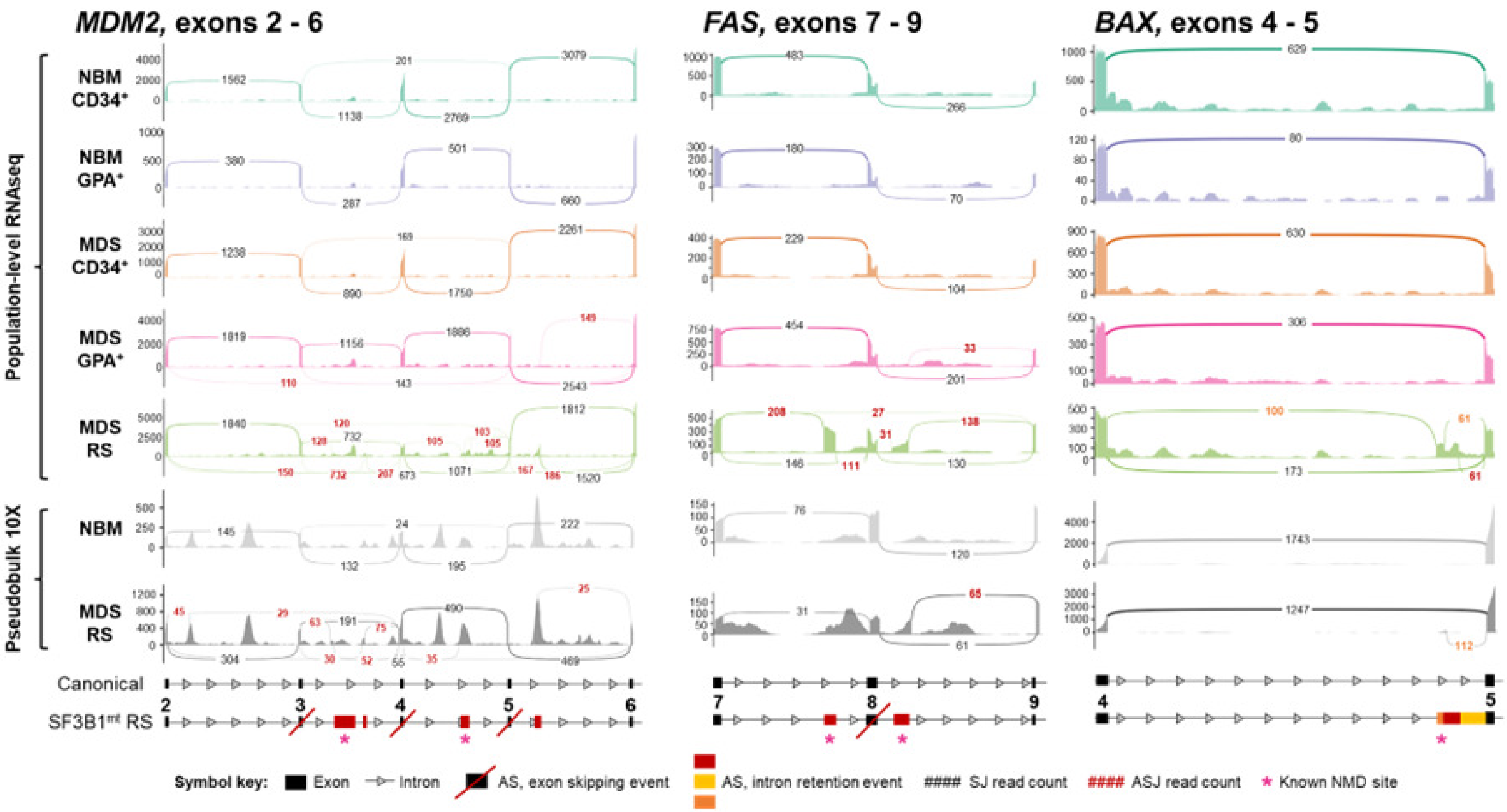

#### Supplemental figure 24

Expanded sashimi plots of population-level RNAseq populations and scRNAseq samples comprising mis-spliced transcript regions of the P53 pathway genes *MDM2*, *FAS* and *BAX*. Canonical splice junction counts (SJ) are noted in black, and alternative SJ counts are noted in red. A full legend is provided below the graph. The asterisks indicate sites corresponding to transcripts known to undergo NMD.
